# Sex-specific and age-related progression of auditory neurophysiological deficits in the *Cln3* mouse model of Batten disease

**DOI:** 10.1101/2025.03.18.643722

**Authors:** Yanya Ding, Jingyu Feng, Viollandi Prifti, Grace Rico, Alexander G. Solorzano, Hayley E. Chang, Edward G. Freedman, John J. Foxe, Kuan Hong Wang

## Abstract

CLN3 disease is a prevalent form of Neuronal Ceroid Lipofuscinosis (NCL) caused by inherited mutations in the *CLN3* gene, with symptoms such as vision loss, language impairment, and cognitive decline. The early onset of visual deficits complicates neurological assessment of brain pathophysiology underlying cognitive decline, while the small number of *CLN3* mutation cases in humans hinders the study of sex differences. Building on our recent progress in assessing auditory neurophysiological changes in CLN3 patients, we developed a parallel approach using electroencephalography arrays in *Cln3* knockout (*Cln3-/-)* mice to investigate the longitudinal progression of auditory processing deficits in both sexes. We employed a duration mismatch negativity (MMN) paradigm, similar to that used in our CLN3 patient studies, to assess the automatic detection of pattern changes in a sequence of stimuli. Wild-type mice of both sexes showed robust duration MMN responses when assessed longitudinally in the same subjects from 3 to 9 months of age. In contrast, female *Cln3-/-* mice developed consistent MMN deficits throughout this age range, while male *Cln3-/-* mice exhibited MMN deficits at younger ages that were mitigated at older ages. Analyses of auditory brainstem responses indicate that MMN abnormalities in *Cln3-/-* mice are not due to peripheral hearing loss. Instead, these deficits originate centrally from sex-specific and age-related changes in auditory evoked potentials elicited by standard and deviant stimuli. Our findings reveal a sex-specific progression of central auditory processing deficits in *Cln3-/-* mice, supporting auditory duration MMN as a translational neurophysiological biomarker for mechanistic studies and therapeutic development.

**Significance Statement:** CLN3 disease is an inherited neurodegenerative disorder with progressive decline in cognitive functioning and verbal abilities. The neuropathophysiological mechanisms underlying this decline remain poorly understood, highlighting the urgent need for objective neurological biomarkers to advance mechanistic insights and therapeutic development. Our identification of central auditory processing and change detection deficits in *Cln3-/-* mice, mirroring findings from our recent studies in CLN3 patients, validates auditory MMN as a translational neurophysiological biomarker bridging pre-clinical and clinical research. Moreover, our discovery of sex-specific, non-linear progression of MMN deficits emphasizes the necessity of developing disease management strategies tailored to each sex. This finding also provides a foundation for investigating both pathogenic and compensatory neural mechanisms to inform the development of individualized treatments.

## Introduction

Neuronal ceroid lipofuscinoses (NCLs), or Batten diseases, are a group of recessively inherited neurodegenerative lysosomal storage disorders (LSDs) that affect children and adolescents (Johnson et al., 2019). CLN3 disease, or juvenile Batten disease, results from mutations in the *ceroid lipofuscinosis neuronal 3 (CLN3)* gene (Consortium, 1995). It is characterized by accumulation of ceroid lipofuscin within lysosomes across various cell types, particularly neurons (Haltia, 2006). In patients, CLN3 disease symptoms typically begin around 4–7 years of age with progressive vision loss, followed by cognitive decline, seizures, motor impairment, language problems, often resulting in premature death around the age of 20 (Mink et al., 2013; Ostergaard, 2016; Tang et al., 2021). Since the early appearance of vision loss followed by cognitive decline complicates the use of standard vision-based neurological evaluations and cognitive assessments, an objective neurophysiological biomarker is needed for monitoring CLN3 disease progression and assessing treatment outcomes.

To develop such a biomarker, previous work in our lab employed electroencephalography (EEG) to measure auditory evoked potentials (AEPs) through the duration mismatch negativity (MMN) paradigm. This approach assesses auditory change detection, a process critical for language comprehension and cognition (Naatanen et al., 2007), both of which are impaired in CLN3 patients (Brima et al., 2024). Since the peripheral auditory system remains relatively intact in CLN3 patients (Adams et al., 2007), AEPs — neural responses time-locked to auditory events — provide valuable insights into the integrity of central auditory processing (Goffin et al., 2014; Modi and Sahin, 2017). Auditory duration MMN is elicited by occasionally presenting deviant stimuli with a different duration within a sequence of standard stimuli (Molholm et al., 2005). Brima and colleagues reported robust auditory duration MMN responses in neurotypical controls, whereas CLN3 patients displayed auditory duration MMN deficits, suggesting impairments in auditory deviant discrimination and auditory sensory memory processes (Brima et al., 2024). While the MMN paradigm is a powerful tool for assessing the integrity of sensory processing (Fitzgerald and Todd, 2020), its use in studying age- and sex-dependent pathophysiological differences in human CLN3 research is constrained by the low prevalence of this genetic disorder (Mirza et al., 2019). Furthermore, the neurophysiological mechanisms underlying auditory MMN dysfunction in CLN3 disease remain unknown.

To establish a common endophenotype with patients for mechanistic studies, we adopted the auditory duration MMN paradigm in the *Cln3-/-* mouse model. Mice were chosen because their AEPs (Modi and Sahin, 2017) and MMN responses (Nagai et al., 2013) are characteristically similar to those in humans, while also providing a controlled genetic system. The *Cln3-/-* mouse model is generated by deleting the first six exons of the *Cln3* gene (Mitchison et al., 1999). It recapitulates the primary pathological hallmark of ceroid lipofuscin accumulation and shows evidence of age-dependent neurodegeneration (Pontikis et al., 2004). *Ex vivo* recordings (Grunewald et al., 2017) and *in vitro* cell culture studies (Kovacs et al., 2006) have identified synaptic transmission dysfunctions in the *Cln3-/-* mouse model associated with behavioral deficits. However, existing animal studies lack neurophysiological measures directly applicable to and comparable with human studies, making it difficult to evaluate the clinical relevance of experimental findings in the *Cln3-/-* mouse model.

In this study, surface EEG electrode arrays, comparable to those used in human studies, were implanted on *Cln3-/-* mice and their wild-type (WT) littermates, enabling longitudinal recordings from the same animals 3-9 months of age using the auditory duration MMN paradigm. *Cln3-/-* mice exhibited prominent auditory duration MMN deficits compared to robust responses in WT mice. To ensure that these deficits were not due to peripheral hearing loss, auditory brainstem responses (ABRs), an objective measure of peripheral auditory processing (Henry, 2002; Zinnamon et al., 2023), were measured. These assessments confirmed that *Cln3-/-* mice showed no peripheral hearing deficits compared to WT mice at the ages tested for MMN. Furthermore, age- and sex-specific deficits in central AEPs that underlie the MMN deficits in *Cln3-/-* mice were revealed. Together, these findings support the use of auditory duration MMN as a translational neurophysiological biomarker for Batten disease.

## Materials and Method

### Animals

*Cln3-/-* mice and their age-matched WT littermates of both sexes on C57BL/6 genetic background were used. Experiments were conducted in accordance with ethical standards for the care and use of animals of the University Committee on Animal Resource (UCAR) at the University of Rochester Medical Center (URMC, NY). Mice were obtained from an in-house breeding colony originally derived from the Jackson Laboratory (B6.129S6-Cln3tm1Nbm/J, JAX:029471, ME). All animals had unlimited access to food and water and were housed in a temperature and humidity-controlled environment with a 12-12h light-dark cycle. No signs of distress were observed in mice during and after EEG or ABR recordings.

### Surgery Procedures

Animals were anesthetized with 5% isoflurane in medical oxygen positioned on a stereotactic frame. They were immediately transferred to a heating pad that maintained temperature around 37°C with 1.5% isoflurane administered through a nose mask at an oxygen flow rate of 2L/min. Then, Meloxicam (1 mg/kg) was injected subcutaneously (SC), and artificial tears (Lubricant Ophthalmic Ointment, Pivetal) were applied to the animal’s eyes. Following preparation, a midline incision was made on the mouse’s scalp. To ensure tight contact of the electrode array to the skull, connective tissue was gently scraped away. In addition, 3% hydrogen peroxide (H_2_O_2_) was applied to remove any membrane residue on the mouse skull. The positioning of the skull was verified by leveling bregma and lambda. Holes for anchoring (right occipital), reference (left occipital) and grounding (frontal) electrode contacts were drilled, followed by immediate insertion of screws. A 32-channel electrode array (H32, NeuroNexus Technologies Inc., MI) was placed on the mouse skull with the middle cross sign positioned at the bregma with a drop of sterilized saline for better electrode attachment to the skull. After the electrode array was correctly placed and excess saline outside the electrode contacts was air dried, Metabond (Parkell C&B, Parkell Inc., NY) was applied to cover the electrode array. Dental acrylic cement was also applied to cover the rest of the exposed skull area and secure the connector to the skull. Incised skin was then sutured, and antibiotic ointment was applied to the area of the sutures and dental cement. Animals were carefully monitored at least once a day for seven consecutive days and analgesics were administered orally or subcutaneously, if necessary.

### Acoustic Stimulation

Acoustic stimuli were generated using RZ6 Multi-I/O processor with Synapse software (Tucker-Davis Technologies, FL) and presented through a speaker (MF1 Multi-Field Magnetic Speaker, Tucker-Davis Technologies, FL), located 10 cm in front of the animal. Sound pressure level (SPL) was modified using programmable attenuators in the TDT system (Tucker-Davis Technologies, FL). Speaker output was calibrated to 80dB SPL at the position of the ears of the mouse in a soundproof recording chamber. Noise burst stimuli were presented at 80dB SPL in blocks of 1000 trials. 850 trials of standard stimuli (50ms in duration) and 150 trials of deviant stimuli (100ms in duration) were played in a pseudo-random manner, with a constant 400ms interstimulus interval (ISI) in each block.

### EEG Data Acquisition and Analysis

Following surgery recovery, animals were habituated in the soundproof recording chamber without acoustic stimulation for at least 3 days. Electrophysiological recording was not performed during habituation. Animals were head-fixed and placed on a rotating disc with a digital head stage connected. Each habituation session lasted 20 minutes. Impedance tests were run during habituation sessions to verify the quality of electrode implantation. Animals with electrode impedance smaller than 30 kΩ or greater than 200 kΩ at a test frequency of 1526 Hz were excluded from the study.

For electrophysiological recording, EEG signals were recorded with an RZ10X Expanded Lux-IO Processor (Tucker-Davis Technologies, FL) at a sampling rate of 1000Hz. Signals were first digitized on the mouse head stage and then filtered during recording by the Synapse software (Tucker-Davis Technologies, FL). Raw data were further processed and analyzed by customized scripts in MATLAB (Mathworks, MA). The applied bandpass filter was 0.1∼100 Hz with the notch filter at 60 Hz.

11 male WT mice, 8 male *Cln3-/-* mice, 6 female WT mice, and 7 female *Cln3-/-* mice were recorded longitudinally from 3 to 9 months of age. Three 5 months old male WT mice and two 5 months old male *Cln3-/-* mice, used for pilot study, were included in addition to longitudinally recorded animals. There were 3 to 5 recording sessions for each animal per age point. Data was first averaged at individual animal level then grouped to compute genotype, age, and sex differences.

Auditory duration MMN was calculated by subtracting AEP to standard stimuli from AEP to deviant stimuli (i.e., MMN = DEV - STD). Auditory duration MMN amplitude was quantified within 140-190ms after stimulus onset, the period during which MMN is maximal in all WT mice. Auditory duration MMN differences were also compared between sex- and age-matched WT and *Cln3-/-* mice across all the electrodes and time points, generating topographical maps of these differences (WT MMN minus *Cln3-/-* MMN) using 20ms time bins. In addition, similar temporospatial analyses were performed for AEPs to examine whether MMN alterations originate from changes in auditory responses to standard stimuli or deviant stimuli.

### ABR Data Acquisition and Analysis

Mice were anesthetized with ketamine (100 mg/kg) and xylazine (10 mg/kg) by intraperitoneal injection (IP). Body temperature was maintained at 37°C by a heating pad. Responses were recorded through subcutaneous needle electrodes placed at the vertex (active), ventrolateral to the left ear (reference), and ventrolateral to the right ear (grounding). Acoustic stimuli were generated by the RZ6 Multi-I/O processor with BioSigRZ Software (Tucker-Davis Technologies, FL) and presented through a speaker (MF1 Multi-Field Magnetic Speaker, Tucker-Davis Technologies, FL), positioned 10 cm from the animal’s ear. Speaker output was calibrated by BioSigRZ software (Tucker-Davis Technologies, FL) with a TDT microphone, placed at the same distance as the mouse’s ear. Decreasing sound pressure levels from 90dB SPL to 10dB SPL in steps of 5dB SPL were employed for ABR stimulation. 512 stimuli, presented at the rate of 21 per second, were recorded for each sound level. 18 male WT mice (4 at 3-month-old, 7 at 5-month-old, 4 at 7-month-old, and 3 at 9-month-old), 20 male *Cln3-/-* mice (6 at 3-month-old, 6 at 5-month-old, 4 at 7-month-old, and 4 at 9-month-old), 16 female WT mice (4 at 3-month-old, 4 at 5-month-old, 4 at 7-month-old, and 4 at 9-month-old), and 18 female *Cln3-/-* mice (5 at 3-month-old, 4 at 5-month-old, 5 at 7-month-old, and 4 at 9-month-old) were used for ABR under anesthesia. The ABR procedure lasted 10 minutes per animal. All animals fully recovered after anesthesia.

ABR data was then processed and analyzed by BioSigRZ Software with a 10ms post-stimulus window. The applied bandpass filter was 300∼30,000 Hz. The notch filter was 60 Hz. The gain was 1 (matched the gain of the preamplifier: Medusa4Z), and the sampling frequency was 12 kHz. Hearing thresholds were determined by visual inspection according to the consensus of four researchers in BioSigRZ Software. Each researcher evaluated raw ABR waveforms individually and determined hearing threshold as the lowest sound level where a waveform could be observed. Researchers were blinded for the age, sex, and genotype of animals they evaluated to avoid bias.

### Statistical Analysis

EEG statistical analyses were performed and plotted using GraphPad Prism 10.2.3 (GraphPad Software, CA) and customized scripts in MATLAB. Mean auditory duration MMN from each channel in the 140–190ms time window was calculated and analyzed using a two-way ANOVA, followed by Fisher’s LSD multiple comparisons for sex-matched WT and *Cln3-/-* mice. Significance was defined as p < 0.05. To explore the spatio-temporal dynamics of the data, two-sided unpaired *t* tests were performed between sex and age-matched WT and Cln3-/-mice across each electrode and time pair (electrode: Ch1-32, time: −50ms to 350ms in 20ms bins). The p-values were corrected for multiple comparisons by controlling the False Discovery Rate (FDR) (Kim et al., 2024)using MATLAB bioinformatics toolbox. Tests were computed for auditory duration MMN, AEP to standard stimuli, and AEP to deviant stimuli separately (i.e., WT MMN vs *Cln3-/-*MMN, WT STD vs *Cln3-/-*STD, WT DEV vs *Cln3-/-*DEV). Results were considered significant if their FDR-corrected p-values were less than 0.05. t-statistics, instead of p-values, were plotted across all the electrodes and time bins using MATLAB to show the direction of changes. Significant results were plotted in dark red and blue, and non-significant results were plotted in light red and blue.

For the statistical analysis of ABR data, two-way ANOVA analysis with Fisher’s LSD multiple comparisons for sex matched WT and *Cln3-/-* mice was performed to determine the effects of age and genotype on hearing thresholds. Significance was defined as p < 0.05.

## Results

### WT mice of both sexes showed robust auditory duration MMN responses

To determine whether auditory duration MMN can be reliably measured in WT mice, a 32-channel electrode array was implanted (Figure 1A) in male and female WT mice, followed by the head-fixed recording of auditory neurophysiological responses using the duration MMN paradigm (Figure 1B). Both male and female WT mice showed clear AEPs in response to standard and deviant stimuli across the skull with left-right symmetry (Extended Data Figure 1-1). The auditory duration MMN in these mice showed a prominent anterior-to-central topographical distribution, with peak responses between 140ms to 190ms post-stimulus onset across all electrodes (Extended Data Figure 1-1). Due to the positioning of the reference and grounding electrodes (Figure 1A), AEPs and auditory duration MMN responses were smaller at the most anterior and posterior electrodes (Extended Data Figure 1-1). Focusing on a centrally located electrode (Ch21), both male and female WT mice displayed robust auditory duration MMN responses from 3 to 9 months of age (Figure 1C and 1D). In addition, topographical maps of AEPs for each electrode, averaged in 20ms increments, showed that AEPs in WT animals propagate from temporal regions near the Auditory Cortex (AC) to central parietal regions, and subsequently to frontal regions (Extended Data Figure 1-2). This propagation pattern aligns with prior literature (Modi and Sahin, 2017) and suggests that EEG recordings in the current study were not dominated by the volume conduction of electrical signals. Overall, WT mice of both sexes showed robust auditory duration MMN responses with minimal age-related changes in AEPs. These findings support the use of EEG recordings with the auditory duration MMN paradigm as a reliable approach for longitudinal studies.

**Figure 1.**
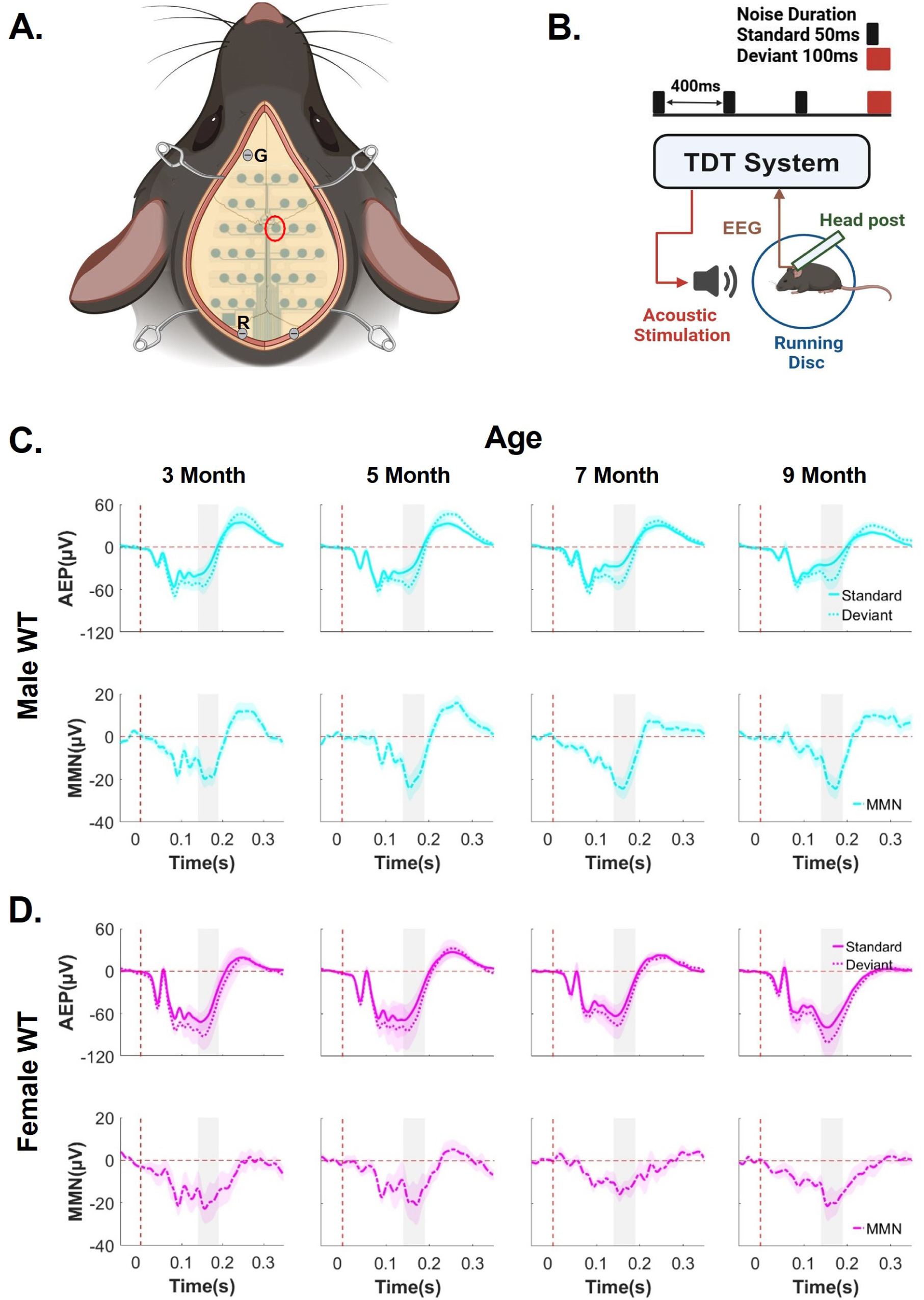
WT mice of both sexes showed robust auditory duration MMN responses from 3 to 9 months of age in longitudinal recordings. ***A,*** Diagram of the 32-channel mouse EEG electrode array implanted on the skull of a mouse. The cross symbol on the array is positioned at the bregma. Screws are inserted for grounding (“G”), reference (“R”), and probe anchorage. ***B***, Diagram shows the setup for recording mouse auditory duration MMN in mice. Animals are head-fixed to a head-post, with EEG and locomotion data recorded using a TDT system. A speaker is placed 10cm in front of the animal. Standard tone: 50ms duration, 850 trials/session; Deviant tone: 100ms duration, 150 trials/session. ***C and D***, Trial and subject averaged AEP and auditory duration MMN waveforms for male (C) and female (D) WT mice aged 3 to 9 months, recorded from a centrally located channel (Ch21, red circle in the channel map). The vertical dashed red line indicates stimulus onset. AEPs in response to standard tones (solid lines) and deviant tones (dotted lines) are presented, with the shaded areas representing the standard error of the mean (SEM). MMN (solid-dotted line), calculated as deviant AEP – standard AEP, are also presented with SEM indicated by shading. The gray rectangle indicates the MMN time window (140–190ms after stimulus onset). Male WT: n = 11 mice for 3-, 7- and 9-month-old groups; n = 14 for the 5-month-old group. Female WT: n = 6 mice for all ages. All animals were recorded longitudinally from 3 to 9 months of age, except three additional male mice recorded only at 5 months of age in pilot studies.

### *Cln3-/-* mice showed sex-specific auditory duration MMN deficits

Next, the effects of *Cln3-/-* genotype on auditory duration MMN responses were examined. As hypothesized, *Cln3-/-* mice showed an auditory duration MMN deficit (Figure 2). Notably, there were sex-specific abnormalities in auditory duration MMN that are associated with disease progression in *Cln3-/-* mice. In male *Cln3-/-* mice, auditory duration MMN responses displayed a dynamic trajectory. At 3 months of age, these mice exhibited strong negativity in the auditory duration MMN response (Figure 2A). However, this response was nearly absent by 5 months of age (Figure 2A). Surprisingly, auditory duration MMN recovered at 7 and 9 months of age (Figure 2A), suggesting the involvement of potential compensatory neural mechanisms in these male mice. In contrast, female *Cln3-/-* mice showed a trend of consistent auditory duration MMN deficits across all ages tested (3 to 9 months) (Figure 2B). Reductions in the magnitude of the auditory duration MMN within the 140-190ms post stimulus time window, the period during which MMN is maximal in all WT mice, were evident at all ages (Figure 2B). In summary, these findings reveal that *Cln3-/-* mice showed sex-specific auditory duration MMN abnormalities, in stark contrast to the consistent and robust responses observed in WT mice.

**Figure 2.**
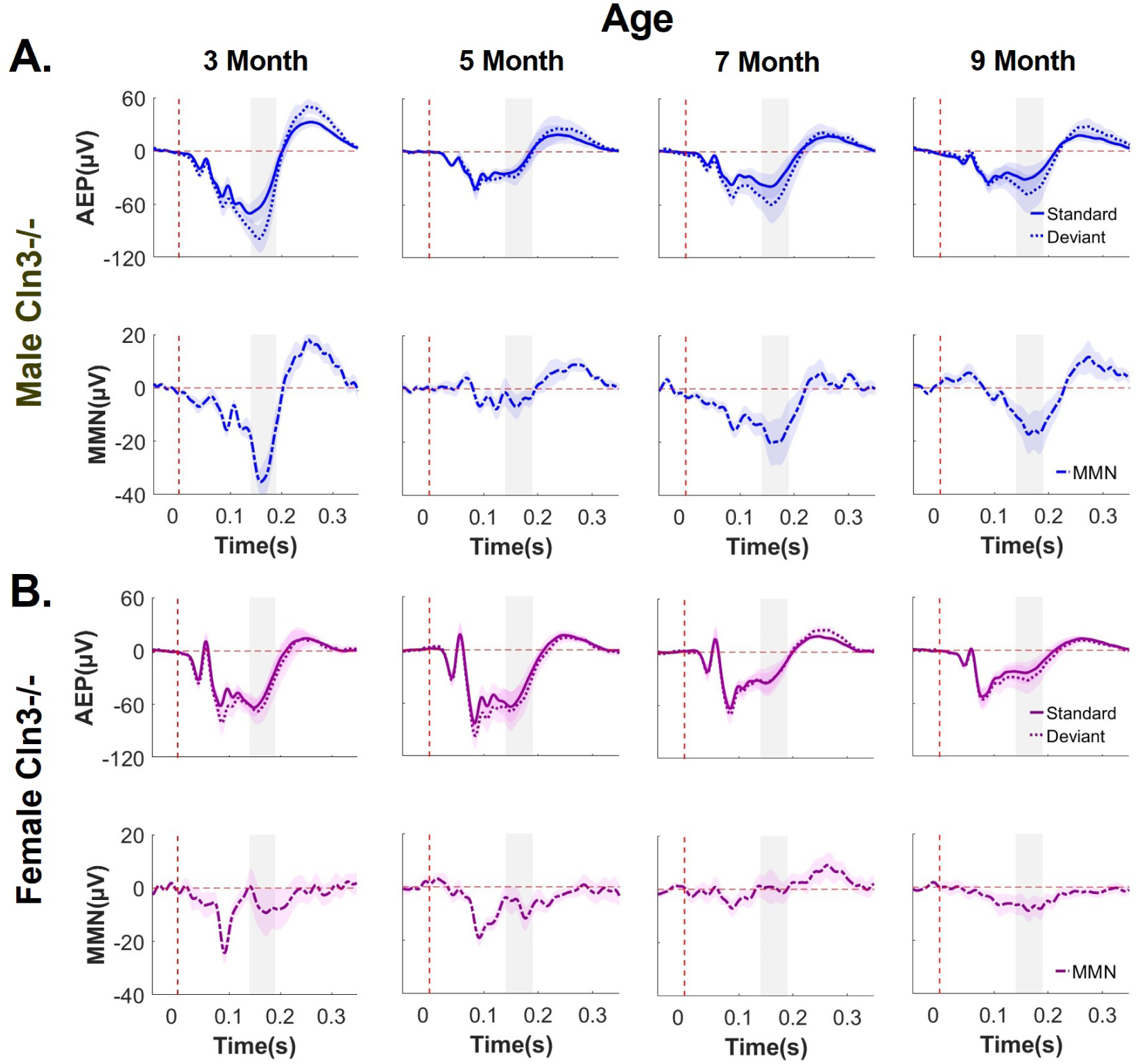
*Cln3*-/- mice showed sex-specific deficits in auditory duration MMN responses. ***A and B*,** Trial and subject averaged AEP and auditory duration MMN waveforms for male (A) and female (B) *Cln3*-/- mice aged 3 to 9 months, recorded from Ch21. The vertical dashed red line indicates stimulus onset. AEPs in response to standard tones (solid lines) and deviant tones (dotted lines) are presented, with the shaded areas representing SEM. MMN (solid-dotted line), calculated as deviant AEP – standard AEP, are also presented with SEM indicated by shading. The gray rectangle indicates the MMN time window (140–190ms after stimulus onset). Male *Cln3-/-* mice showed a clear auditory duration MMN at 3 months of age, a diminished MMN at 5 months, and a subsequent recovery at 7 and 9 months (A), whereas female *Cln3-/-* mice showed diminished MMN across all ages (B). Male *Cln3-/-*: n = 8 mice for 3-, 7- and 9-month-old groups; n = 10 for the 5-month-old group. Female *Cln3-/-*: n = 7 mice for all ages. All animals were recorded longitudinally from 3 to 9 months of age, except two additional male mice recorded only at 5 months of age in pilot studies.

### MMN deficits emerged in younger *Cln3-/-* males were mitigated at older ages

To further examine the effects of *Cln3* mutations on central auditory processing, the spatio-temporal patterns of auditory duration MMN responses in *Cln3-/-* mice were compared with those of sex- and age-matched WT mice. In male *Cln3-/-* mice, amplitudes of auditory duration MMN at a central electrode (Ch21) were elevated at 3 months of age, reduced at 5 months, and subsequently restored to approximately the WT level at 7 and 9 months (Figure 3A). This age-dependent up-and-down modulation of auditory duration MMN magnitudes was consistently observed across electrodes at various spatial locations (Extended Data Figure 3-1 to 3-5). For statistical analysis, the mean auditory duration MMN was calculated by averaging the response over the 140–190ms time window and tested the effects of age and genotype (Figure 3B). Although there was no significant main effect of age (F (3,73) = 1.818, p = 0.1515, η_p_² = 0.06366, two-way ANOVA) or genotype (F(1,73) = 0.6395, p = 0.4265, η_p_² = 0.007467, two-way ANOVA), the interaction effect between age and genotype showed a trend toward significance (F(3,73) = 2.639, p = 0.0559, η_p_² = 0.09244, two-way ANOVA). This likely reflects the non-linear changes in MMN across age, with 5-month-old male Cln3-/- mice showing significantly reduced mean auditory duration MMN compared to age-matched WT mice (p = 0.01, Fisher’s LSD multiple comparisons test) (Figure 3B).

**Figure 3.**
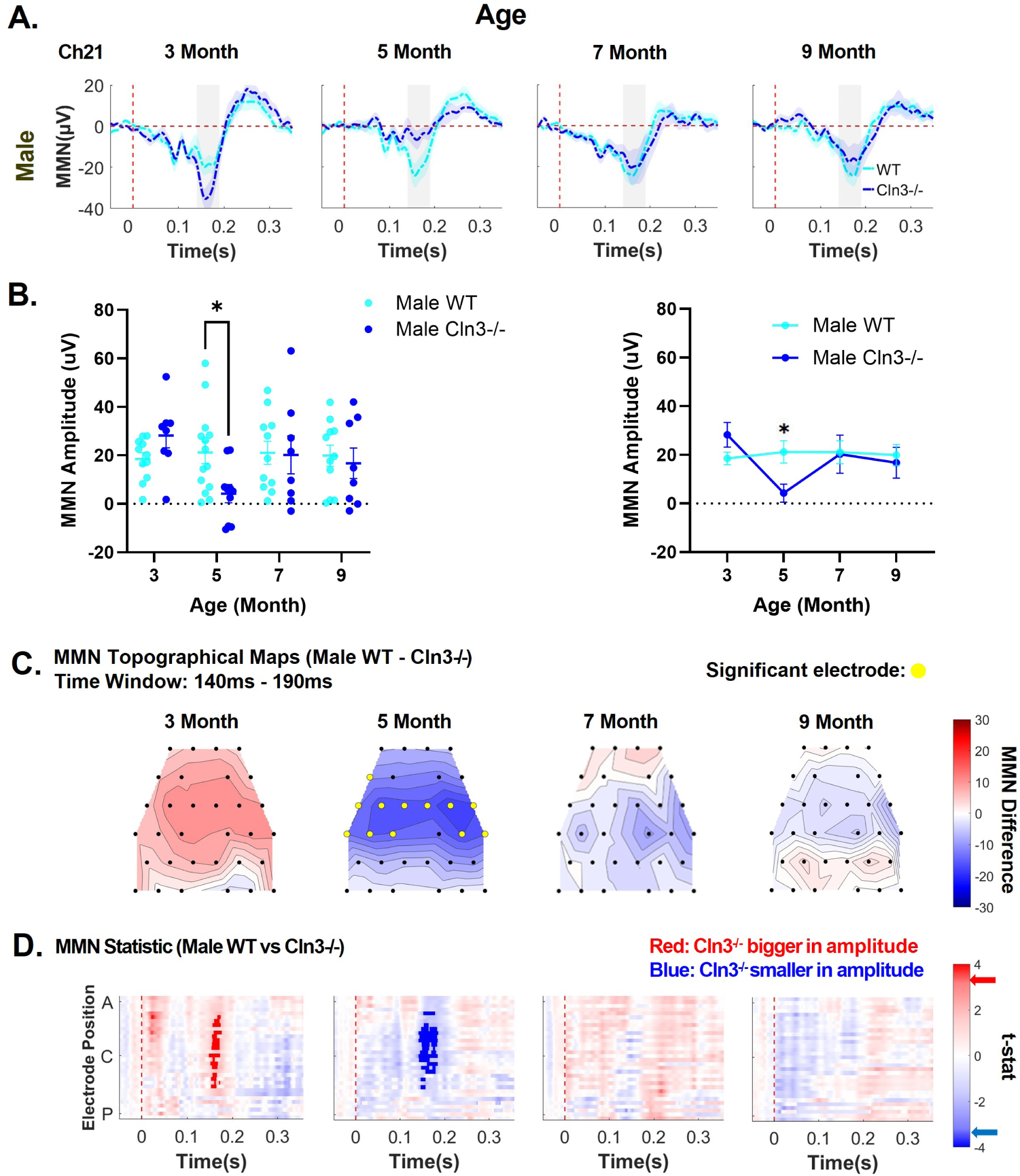
Male *Cln3-/-* mice showed an initially greater auditory duration MMN response compared to WT mice, followed by a clear reduction and subsequent recovery. ***A,*** Plots comparing trial- and subject-averaged auditory duration MMN waveforms for male WT mice (light blue) and male *Cln3-/-* mice (dark blue) at Ch21. The vertical dashed red line indicates stimulus onset. The gray rectangle indicates the MMN time window (140–190ms after stimulus onset). ***B*,** Mean MMN amplitude within the 140-190ms time window at Ch21 for male WT and *Cln3-/-* mice. The left scatter plot shows individual data points for each animal, whereas the right line plot shows mean ± SEM. Two-way ANOVA with Fisher’s LSD comparison showed a significant reduction in the mean auditory duration MMN in 5-month-old *Cln3-/-* mice compared to WT mice (p = 0.01). ***C*,** Topographical maps of the mean auditory duration MMN difference within the 140-190ms time window between male WT and male *Cln3-/-* mice (i.e., WT MMN – *Cln3-/-* MMN). Red indicates regions where *Cln3-/-* mice showed larger amplitudes (further away from zero compared to WT mice), while blue indicates regions where *Cln3-/-* mice showed smaller amplitudes (closer to zero compared to WT mice). Black dots indicate electrode positions, while yellow dots highlight electrodes showing significant MMN differences. ***D*,** Statistical differences in auditory duration MMN between male WT and *Cln3-/-* mice were displayed across all electrodes and the entire trial duration (32 electrodes × 400ms). Results from *t*-tests were corrected for multiple comparisons using the false discovery rate (FDR) method. Significance was determined when the FDR-adjusted p-value was less than 0.05 (two-tailed). Electrode positions over the mouse skull are indicated from A (anterior) to C (center) and P (posterior). Red indicates electrodes and time bins (in 20ms) where *Cln3-/-* mice showed larger amplitudes (further away from zero compared to WT mice), while blue indicates electrodes and time bins where *Cln3-/-* mice showed smaller amplitudes (closer to zero compared to WT mice). Significant t-stat values are represented by dark red and dark blue, based on the FDR-corrected p-value. Red and blue arrows on the color bar indicate significant t-stat value. Male WT: n = 11 mice for 3-, 7- and 9-month-old groups; n = 14 for the 5-month-old group. Male *Cln3-/-*: n = 8 mice for 3-, 7- and 9-month-old groups; n = 10 for the 5-month-old group. Male *Cln3-/-* mice initially exhibited an enhanced auditory duration MMN response within the 140-190ms time window at 3 months of age, followed by a clear reduction at 5 months, and subsequently showed recovery at 7 and 9 months of age.

Next, the spatial distribution of MMN differences between WT and *Cln3-/-* male mice was examined using topographical maps. While male WT mice consistently showed robust auditory duration MMN across multiple electrodes and ages, male *Cln3-/-* mice displayed age-dependent variations in the spatial distribution of MMN (Extended Data Figure 3-6). To better visualize genotype differences, topographical maps were generated by subtracting the MMN of male *Cln3-/-* mice (averaged over the 140–190ms time window) from that of male WT mice. At 3 months of age, male *Cln3-/-* mice showed greater MMN amplitudes compared to WT mice, with the most pronounced differences observed at central electrodes (Figure 3C). By 5 months, the genotype difference reversed, with male *Cln3-/-* mice exhibiting reduced MMN amplitudes relative to WT mice, again showing the largest differences at central electrodes (Figure 3C). At 7 and 9 months, little to no genotype difference in the spatial distribution of MMN was observed, suggesting an age-dependent compensation of the initial differences (Figure 3C).

To test the statistical significance of genotype differences in the temporal and spatial distributions of MMN in male mice, pairwise *t*-tests were performed across all electrodes and time points and corrected for multiple comparisons by controlling the FDR (see Methods). At 3 months of age, male *Cln3-/-* mice showed a significant increase in MMN magnitude compared to WT mice at central electrodes, particularly within the 150– 170ms post-stimulus time window (Figure 3D). At 5 months, male Cln3-/- mice exhibited significantly reduced MMN magnitudes relative to WT mice, with the largest differences observed at central electrodes from 140–180ms post-stimulus onset (Figure 3D). By 7 and 9 months of age, there were no significant differences in auditory duration MMN between male WT and Cln3-/- mice. In summary, while MMN magnitudes remained stable across age in male WT mice, male *Cln3-/-* mice showed age-dependent fluctuations in MMN amplitude, with an initial increase followed by a reduction and eventual restoration to levels comparable to WT mice at older ages.

### A trend of MMN deficits across age in Cln3-/- females

Next, genotype differences in the spatio-temporal patterns of auditory duration MMN response in female mice were examined. The data showed that MMN magnitude in the 140-190ms post-stimulus time window was consistently reduced in *Cln3-/-* mice compared to WT mice across all ages at Ch21 (Figure 4A). This reduction in MMN magnitude was also observed consistently across other electrodes (Extended Figures 4-1 to 4-5). Statistical analysis at Ch21 revealed a significant genotype effect, with *Cln3-/-* mice showing reduced mean auditory duration MMN amplitudes compared to WT mice (F(1,44) = 9.848, p = 0.0030, η_p_² = 0.1756, two-way ANOVA) (Figure 4B). However, there were no significant effects of age (F(3,44) = 0.7146, p = 0.5485, η_p_² = 0.03822, two-way ANOVA) or genotype-by-age interaction (F(3,44) = 0.02228, p = 0.9954, η_p_² = 0.001192, two-way ANOVA) (Figure 4B).

**Figure 4.**
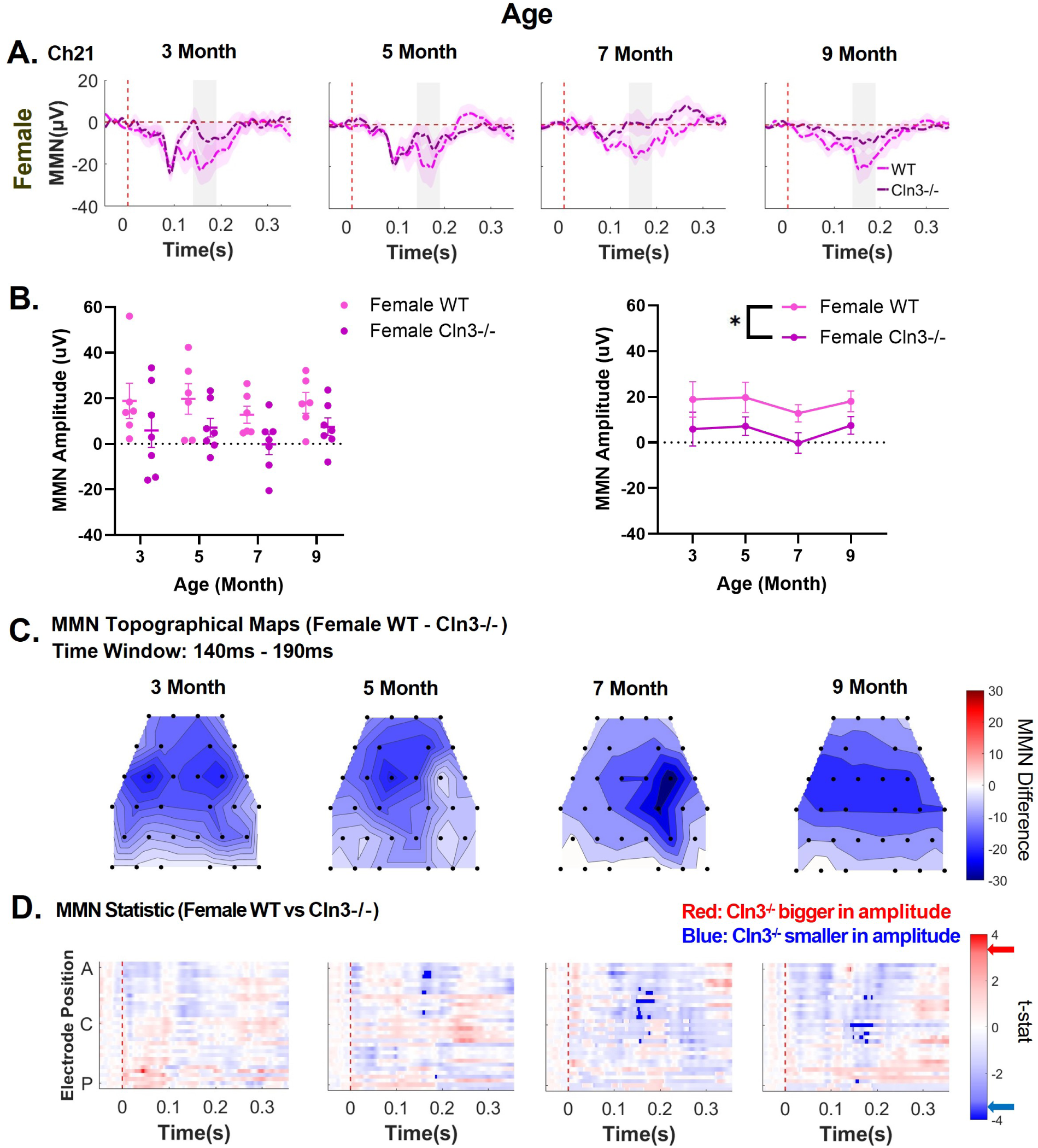
Female *Cln3-/-* mice showed consistent auditory duration MMN deficits from 3 to 9 months of age. ***A***, Plots comparing trial- and subject-averaged auditory duration MMN waveforms for female WT mice (light pink) and female *Cln3-/-* mice (dark pink) at Ch21. The vertical dashed red line indicates stimulus onset. The gray rectangle indicates the MMN time window (140–190ms after stimulus onset). ***B*,** Mean MMN amplitude within the 140-190ms time window at Ch21 for female WT and *Cln3-/-* mice. The left scatter plot shows individual data points for each animal, whereas the right line plot shows mean ± SEM. Two-way ANOVA showed a significant genotype main effect (p= 0.0030), with a reduction in the mean auditory duration MMN in female *Cln3-/-* mice compared to female WT mice. ***C***, Topographical maps of the mean auditory duration MMN difference within the 140-190ms time window between female WT and female *Cln3-/-* mice (i.e., WT MMN – *Cln3-/-* MMN). Red indicates regions where *Cln3-/-* mice showed larger amplitudes (further away from zero compared to WT mice), while blue indicates regions where *Cln3-/-* mice showed smaller amplitudes (closer to zero compared to WT mice). Black dots indicate electrode positions. ***D***, Statistical differences in auditory duration MMN between female WT and *Cln3-/-* mice were displayed across all electrodes and the entire trial duration (32 electrodes × 400ms). Results from *t*-tests were corrected for multiple comparisons using the FDR method. Significance was determined when the FDR-adjusted p-value was less than 0.05 (two-tailed). Electrode positions over the mouse skull are indicated from A (anterior) to C (center) and P (posterior). Red indicates electrodes and time bins (in 20ms) where *Cln3-/-* mice showed larger amplitudes (further away from zero compared to WT mice), while blue indicates electrodes and time bins where *Cln3-/-* mice showed smaller amplitudes (closer to zero compared to WT mice). Significant *t*-stat values are represented by dark red and dark blue, based on the FDR-corrected p-value. Red and blue arrows on the color bar indicate significant t-stat value. Female WT: n = 6 mice for all ages. Female *Cln3-/-*: n = 7 mice for all ages. Female *Cln3-/-* mice showed a trend of consistent auditory duration MMN deficits within the 140-190ms time window at central to anterior electrodes, with greater significance at later ages.

Topographical maps of MMN genotype differences further confirmed that female *Cln3-/-* mice showed reduced MMN magnitudes compared to the WT in the 140-190ms time window across multiple electrodes and age groups (Figure 4C). While female WT mice consistently showed robust auditory duration MMN across multiple electrodes and ages, female *Cln3-/-* mice displayed an overall lower magnitude in the spatial distribution of MMN (Extended Data Figure 4-6). Detailed analyses of genotype differences at individual electrodes and time points also revealed significant MMN reductions in female *Cln3-/-* mice at certain electrodes and time points (Figure 4D). In summary, female *Cln3-/-* mice showed a trend of auditory duration MMN deficits across ages.

### *Cln3-/-* and WT mice showed similar peripheral hearing from 3 to 9 months of age

To rule out the possibility that auditory duration MMN deficits originate from peripheral hearing loss, click ABR recording was performed to test hearing thresholds in WT and *Cln3-/-* mice of both sexes at the ages corresponding to the EEG recordings (Figure 5A and 5B). Hearing thresholds in all animals were below the testing sound level (i.e., 80dB SPL) of our EEG stimuli (Figure 5C and 5D). In male mice, there were no significant effects of genotype (F(1,30) = 0.3363, p = 0.5663, η_p_² = 0.005533, two-way ANOVA) or genotype-by-age interaction (F(3,30) = 0.3922, p-value = 0.7595, η_p_² = 0.01936, two-way ANOVA) (Figure 5C). However, as expected, a significant age effect was observed (F(3,30) = 9.571, p = 0.0001, η_p_² = 0.4725, two-way ANOVA), reflecting a gradual increase in hearing thresholds with age (Figure 5C). Similarly, in female mice, there was a significant effect of age (F(3,26) = 11.17, p < 0.0001, η_p_² = 0.4946, two-way ANOVA), but no significant effects of genotype (F(1,26) = 3.662, p = 0.0668, η_p_² = 0.05402, two-way ANOVA) or genotype-by-age interaction (F(3,26) = 1.681, p = 0.1955, η_p_² = 0.07441, two-way ANOVA) (Figure 5D).

**Figure 5.**
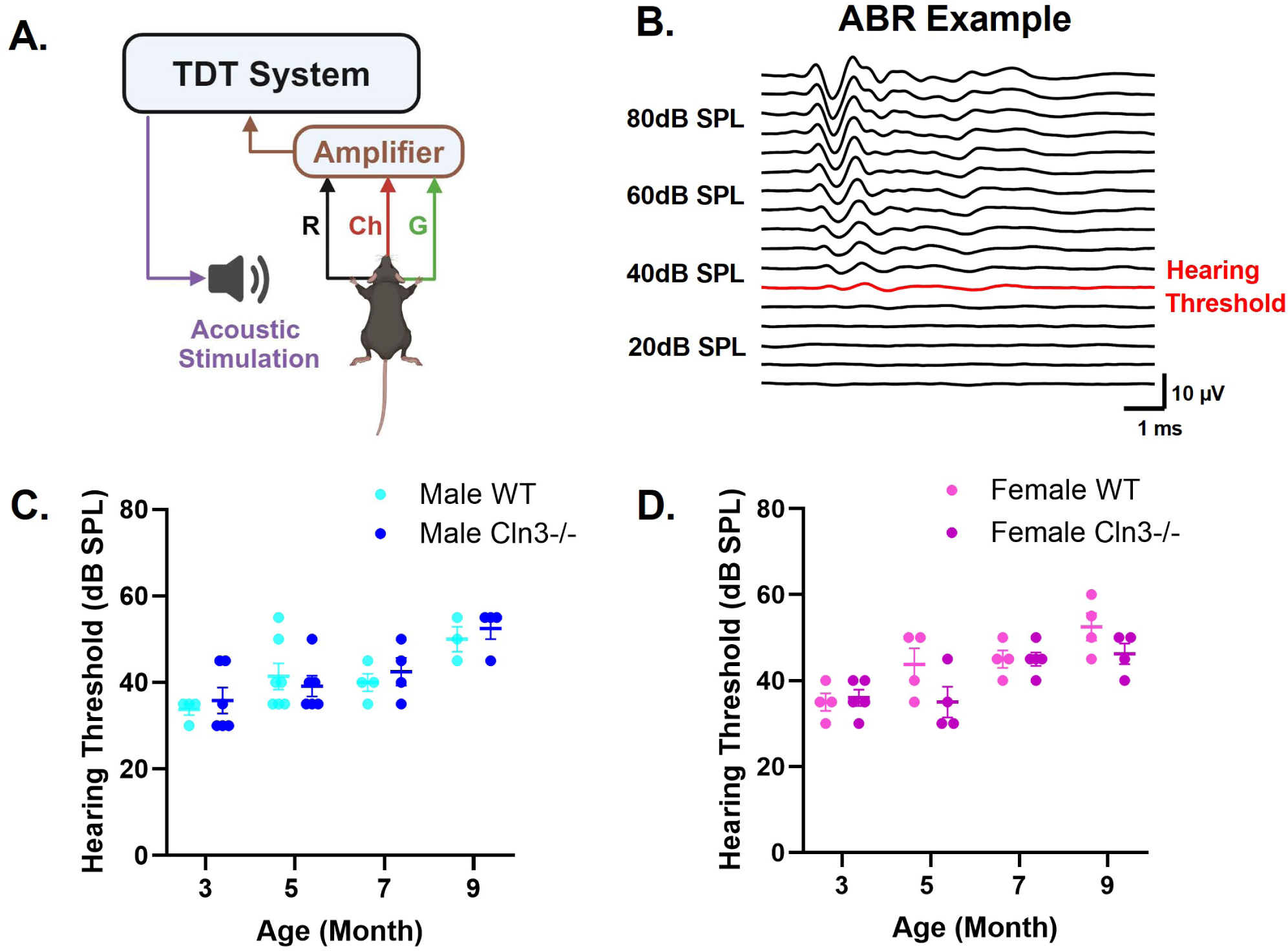
*Cln3-/-* and WT mice showed similar peripheral hearing from 3 to 9 months of age. ***A***, Diagram shows the click ABR recording set up. The TDT system is connected to a speaker for stimulus delivery. ABRs are recorded with three needle electrodes: black for reference (R), green for grounding (G), and red for active recording channel (Ch). The signals are amplified by an amplifier and processed by the TDT system. ***B,*** The ABR waveforms of an example animal are shown, ranging from 10dB to 90dB. The hearing threshold for this animal is indicated in red. ***C***, Click ABR hearing thresholds for male WT (light blue) and male *Cln3-/-* mice (dark blue). Male *Cln3-/-* mice (n = 6 at 3-months, n = 6 at 5 months, n = 4 at 7 months, n = 4 for 9 months) showed no significant difference in click ABR hearing thresholds compared to age-matched male WT mice (n = 4 at 3 months, n = 7 at 5 months, n = 4 at 7 months, n = 3 at 9 months). ***D***, Click ABR hearing thresholds for female WT (light pink) and female *Cln3-/-* mice (dark pink). Female *Cln3-/-* mice (n = 5 at 3- and 7 months, n = 4 at 5 and 9 months) showed no significant difference in click ABR hearing thresholds compared to age-matched female WT mice (n = 4 for all ages). From 3 to 9 months of age, the hearing thresholds of all WT and *Cln3-/-* mice are below the testing sound level (i.e., 80dB SPL) used in the auditory duration MMN paradigm.

The observed age-related differences in hearing thresholds for both sexes were consistent with the trend of age-related hearing loss (AHL) reported in the C57BL/6J mouse strain(Ison et al., 2007). Previous studies have documented complete hearing loss in this strain between 12 and 15 months of age (Someya et al., 2009). Additional recordings in 13-month-old mice confirmed that animals with complete hearing loss had no identifiable ABR or AEP waveforms (Extended Data Figure 5-1). In conclusion, WT and *Cln3-/-* mice showed similar peripheral hearing thresholds at the ages corresponding to EEG recordings, indicating that auditory duration MMN deficits in *Cln3-/-* mice arise from central dysfunction rather than peripheral hearing loss.

### Male *Cln3-/-* mice showed age-dependent up-and-down modulation of AEPs

To investigate whether auditory duration MMN deficits in *Cln3-/-* mice originate from changes in auditory responses to standard stimuli, deviant stimuli, or both, AEPs between sex- and age-matched WT and *Cln3-/-* mice were compared. Standard AEPs in 3-month-old male *Cln3-/-* mice showed increased amplitude in several anterior-to-central electrodes within the typical MMN window (140-190ms) compared to male WT mice (Figure 6A). Deviant AEPs in these *Cln3-/-* mice also exhibited enhanced amplitude compared to WT mice, spanning more electrodes and time points than the standard AEPs (Figure 6B). However, at 5 months of age, both standard and deviant AEPs in *Cln3-/-* mice were significantly reduced compared to WT mice, with reductions in deviant AEPs affecting more central electrodes and time points within the MMN window. By 7 and 9 months of age, both standard and deviant AEPs in male *Cln3-/-* mice returned to levels comparable to those of WT mice within the MMN time window (Figure 6). Topographical maps of standard and deviant AEPs in male mice, averaged in 20ms time bins, further illustrate these spatiotemporal changes (Extended Data Figure 6-1 and 6-2). In summary, male *Cln3-/-* mice showed age-dependent up-and-down modulation of AEP amplitudes, with more dramatic changes in deviant AEPs contributing to the genotype differences in MMN.

**Figure 6.**
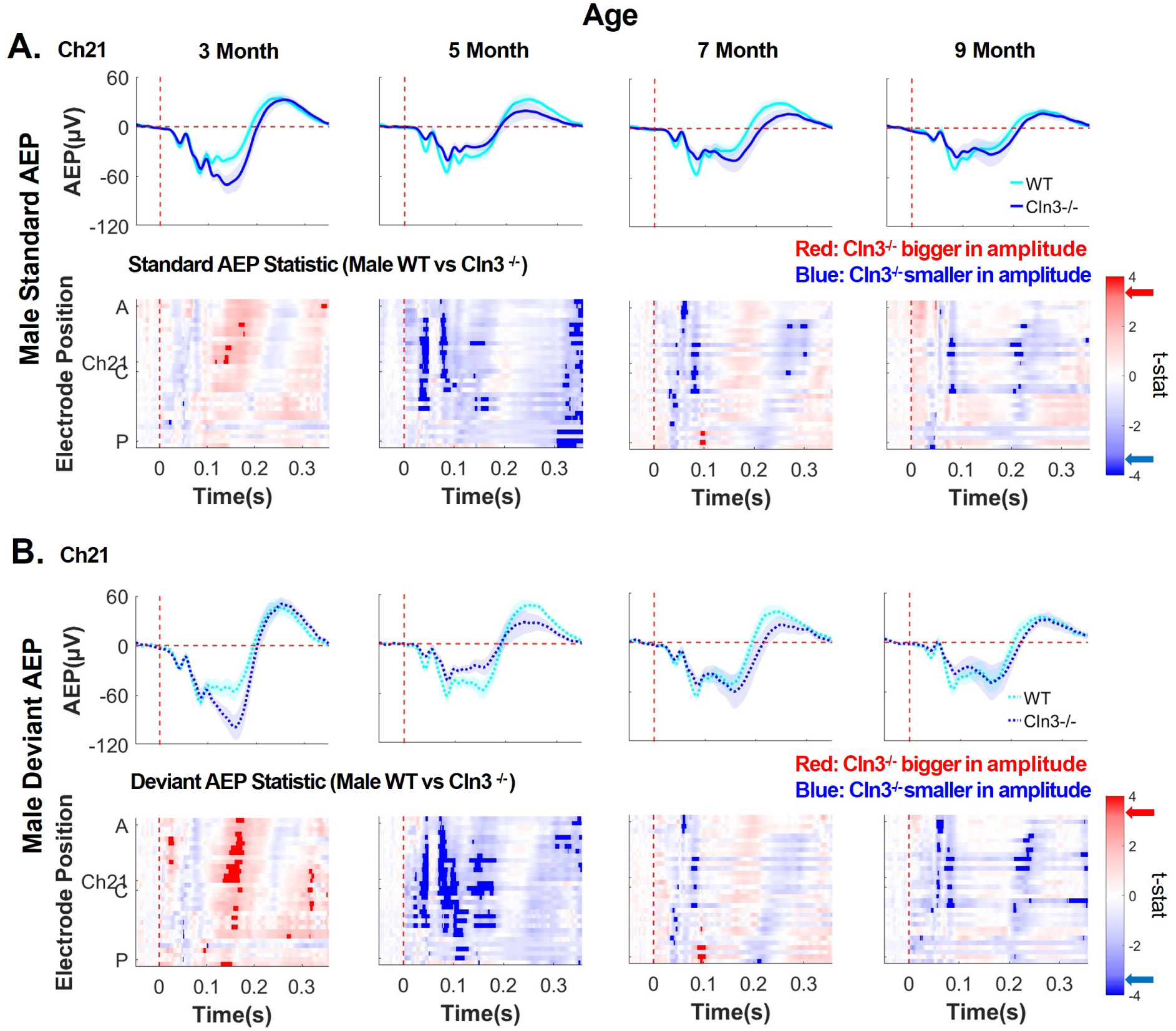
Male *Cln3-/-* mice initially exhibited a greater AEP response compared to WT mice, followed by a decline and subsequent recovery. ***A,*** top row, trial- and subject-averaged standard AEP waveforms for male WT mice (light blue) and male *Cln3-/-* mice (dark blue) at Ch21. The vertical dashed red line indicates stimulus onset. Bottom row, statistical differences in standard AEP between male WT and *Cln3-/-* mice were displayed across all electrodes and the entire trial duration (32 electrodes × 400ms). Results from *t*-tests were corrected for multiple comparisons using the FDR method. Significance was determined when the FDR-adjusted p-value was less than 0.05 (two-tailed). Electrode positions over the mouse skull are indicated from A (anterior) to C (center) and P (posterior). Red indicates electrodes and time bins (in 20ms) where *Cln3-/-* mice showed larger amplitudes (further away from zero compared to WT mice), while blue indicates electrodes and time bins where *Cln3-/-* mice showed smaller amplitudes (closer to zero compared to WT mice). Significant *t*-stat values are represented by dark red and dark blue, based on the FDR-corrected p-value. Red and blue arrows on the color bar indicate significant t-stat value. ***B*,** Deviant AEP waveforms at Ch21 (top row) and *t*-stat values across all electrodes and the entire trial duration (bottom row). Male WT: n = 11 mice for 3-, 7- and 9-month-old groups; n = 14 for the 5-month-old group. Male*Cln3-/-*: n = 8 mice for 3-, 7- and 9-month-old groups; n = 10 for the 5-month-old group. Male *Cln3-/-* mice initially exhibited an enhanced auditory AEP response within the 140-190 ms MMN time window at 3 months of age, followed by a reduction at 5 months, and subsequently showed recovery at 7 and 9 months of age. Deviant AEP changes impacted more central electrodes and timepoints within the MMN window compared to standard AEPs.

### Female Cln3-/- mice showed a progressive decline in AEPs

Age-related changes in AEPs in female mice were also examined. While female WT mice maintained consistent AEPs from 3 to 9 months of age, female *Cln3-/-* mice showed a gradual decline in AEP magnitudes (Figure 7). Both standard and deviant AEPs within the typical MMN time window (140–190ms post-stimulus onset) were reduced in female *Cln3-/-* mice compared to age-matched WT mice, with the most pronounced differences observed at 9 months of age in anterior-to-central electrodes (Figure 7A and B). Topographical maps of standard and deviant AEPs, averaged in 20ms time bins, further illustrate these spatiotemporal changes in female mice (Extended Data Figure 7-1 and 7-2). In summary, female *Cln3-/-* mice demonstrated a progressive decline in both standard and deviant AEPs, contributing to a consistent trend of auditory duration MMN deficits.

**Figure 7.**
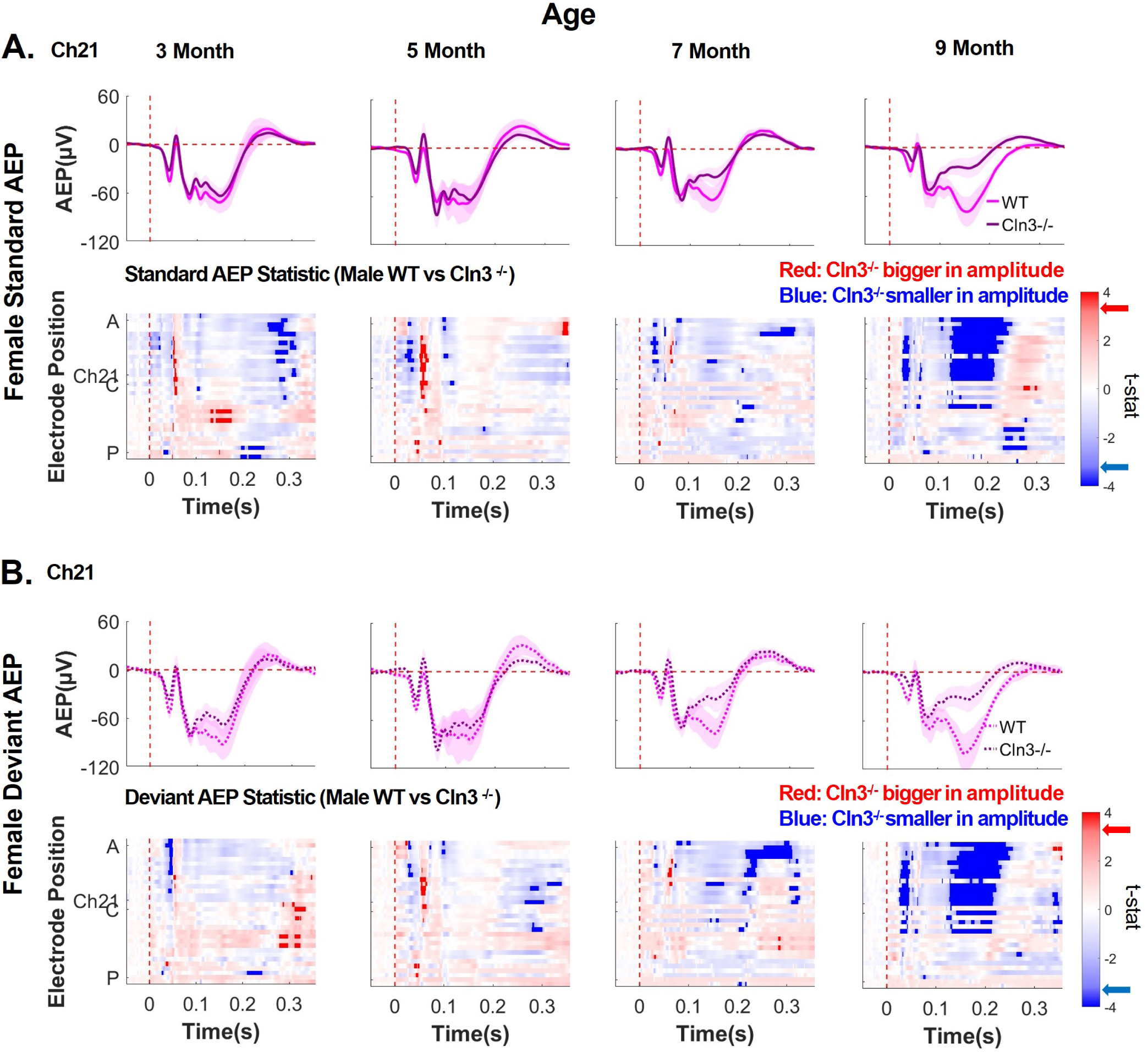
Female *Cln3-/-* mice showed a progressive decline in AEPs compared to WT mice. ***A***, top row, trial- and subject-averaged standard AEP waveforms for female WT mice (light pink) and *Cln3-/-* mice (dark pink) at Ch21. The vertical dashed red line indicates stimulus onset. Bottom row, statistical differences in standard AEP between female WT and *Cln3-/-* mice were displayed across all electrodes and the entire trial duration (32 electrodes × 400ms). Results from *t*-tests were corrected for multiple comparisons using the FDR method. Significance was determined when the FDR-adjusted p-value was less than 0.05 (two-tailed). Electrode positions over the mouse skull are indicated from A (anterior) to C (center) and P (posterior). Red indicates electrodes and time bins (in 20ms) where *Cln3-/-* mice showed larger amplitudes (further away from zero comparing to WT mice), while blue indicates electrodes and time bins where *Cln3-/-* mice showed smaller amplitudes (closer to zero comparing to WT mice). Significant *t*-stat values are represented by dark red and dark blue, based on the FDR-corrected p-value. Red and blue arrows on the color bar indicate significant t-stat value. ***B***, Deviant AEP waveforms at Ch21 (top row) and *t*-stat values across all electrodes and the entire trial duration (bottom row). Female WT: n = 6 mice for all ages. Female *Cln3-/-*: n = 7 mice for all ages. AEPs within the typical MMN time window (140–190ms) progressively decreased with age in female *Cln3-/-* mice. By 9 months of age, female *Cln3-/-* mice showed significantly reduced standard and deviant AEPs at anterior to central electrodes.

### Sex-specific and age-dependent alterations of auditory duration MMN and AEPs in *Cln3-/-* mice

To summarize the sex-specific and age-dependent alterations in auditory duration MMN and AEPs in the *Cln3-/-* mouse model, the area under the curve (AUC) of averaged MMN and AEP waveforms was computed for each group at Ch21, spanning 50ms before stimuli onset to 350ms post stimuli onset. The ratio of AUC values between *Cln3-/-* and WT mice was then calculated. Male *Cln3-/-* mice initially exhibited a greater response than WT mice (ratio > 1), followed by a reduced response (ratio < 1), and subsequently showed recovery in both auditory duration MMN and AEPs from 3 to 9 months of age (Figure 8). In contrast to the non-linear up-and-down alterations in male *Cln3-/-* mice, female *Cln3-/-* mice showed a progressive, age-dependent decline in both AEPs and MMN over the same period (Figure 8). Overall, alterations in auditory duration MMN closely mirrored changes in AEPs for both male and female *Cln3-/-* mice.

**Figure 8.**
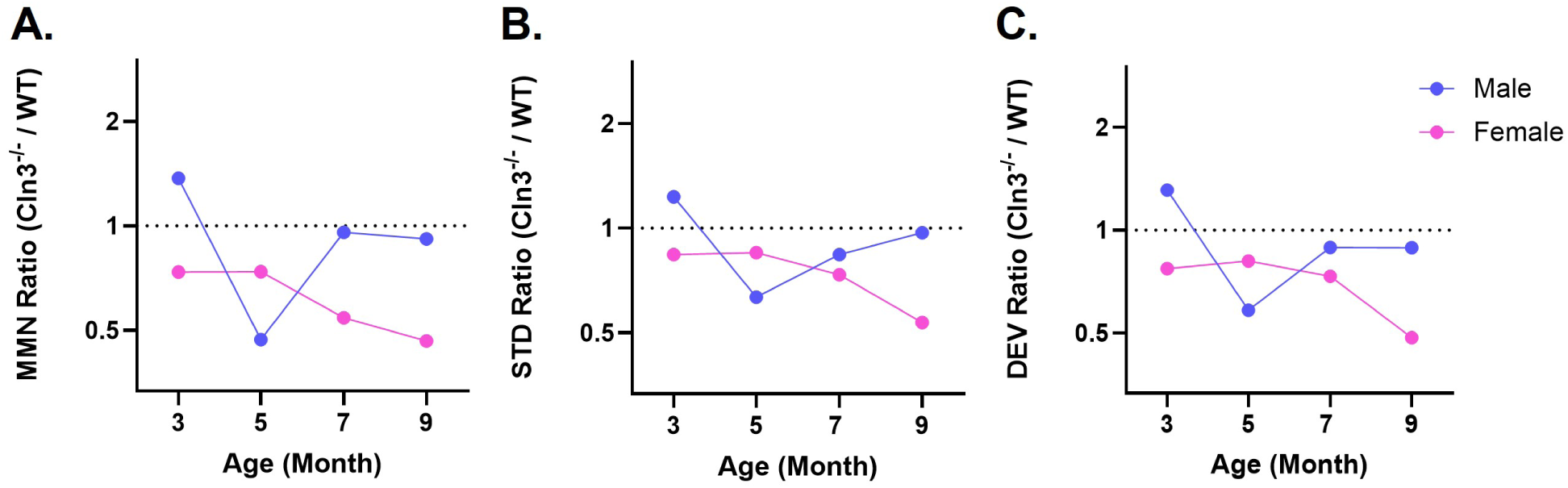
Auditory duration MMN abnormalities originate from parallel changes in AEPs. ***A-C***, Summary trends for auditory duration MMN (A), standard AEP (B), and deviant AEP (C) changes in *Cln3-/-* mice compared to WT mice across ages. Ratios are computed as the group-averaged area under the curve (AUC) of *Cln3-/-* mice divided by that of WT mice from Ch21 across all ages (i.e., *Cln3-/-* AUC / WT AUC). A ratio greater than 1 indicates larger amplitudes in *Cln3-/-* mice, while a ratio less than 1 indicates smaller amplitudes in *Cln3-/-* mice. ***A***, Compared to male WT mice, male *Cln3-/-* mice exhibited larger auditory duration MMN amplitudes at 3 months of age, smaller amplitudes at 5 months, and similar amplitudes at 7 and 9 months. In contrast, female *Cln3-/-* mice showed consistently smaller auditory duration MMN amplitudes compared to female WT mice across all ages, with greater reductions observed at later ages. ***B***, Compared to male WT mice, male *Cln3-/-* mice exhibited an initially larger amplitude of standard AEP, followed by a reduction and gradual recovery with age. In contrast, female *Cln3-/-* mice showed an overall reduction in AEPs, with greater decreases observed at later ages. ***C,*** Compared to male WT mice, male *Cln3-/-* mice exhibited an initially larger amplitude of deviant AEP, followed by a reduction and gradual recovery with age. In contrast, female *Cln3-/-* mice showed an overall reduction in AEPs, with greater decreases observed at later ages.

## Discussion

The current study identified central auditory processing dysfunctions in the *Cln3-/-* mouse model. While WT mice for both sexes showed consistent AEPs with robust auditory duration MMN response from 3 to 9 months of age, *Cln3-/-* mice displayed sex- and age-specific deficits of auditory duration MMN, paralleling changes in AEPs. In male *Cln3-/-* mice, amplitudes of auditory duration MMN and AEPs were initially elevated at 3 months, reduced at 5 months, and subsequently restored to levels comparable to WT mice at 7 and 9 months. In contrast, female *Cln3-/-* mice exhibited a progressive decline in AEPs with a consistent trend of auditory duration MMN deficits from 3 to 9 months of age. Prior studies of *Cln3* mouse models have largely focused on middle-aged to older animals (6–24 months), where storage body accumulation (Katz et al., 2008) and glial activation (Langin et al., 2020) are more pronounced. Although sex differences in behavioral and motor outcomes have been reported in *Cln3* mouse models (Kovacs and Pearce, 2015), neurophysiological responses remain underexplored, particularly in female mice, as most studies have prioritized male subjects (McShane and Mole, 2022). This study provides the first demonstration of age-related and sex-specific central auditory processing abnormalities in the *Cln3-/-* mouse model of Batten disease. It emphasizes the importance of systematically examining subjects across different ages and both sexes to fully understand the physiological consequences of CLN3 mutations and to inform the development of targeted disease management and treatment strategies.

Findings of the current study validate the utility of the auditory duration MMN paradigm in murine models for longitudinal assessment of sensory processing deficits. While MMN paradigms have been used in some rodent models of brain disorders, for example, to capture deficits potentially related to schizophrenia (Lee et al., 2018; Jodo et al., 2019), their use for long-term tracking of neurophysiological deficits remains limited. Here, this study demonstrates that auditory neurophysiological responses in WT mice are consistent across six months of adult life, confirming the reliability of the auditory duration MMN paradigm for assessing auditory sensory memory and deviance detection over time(Garrido et al., 2009). This paradigm can also serve as a functional outcome measure to evaluate the efficacy of therapeutic strategies, such as optogenetic manipulations, pharmaceuticals, and gene therapies. For example, AAV9-mediated human CLN3 (hCLN3) gene therapy has shown effects in restoring CLN3 protein expression (Bosch et al., 2016) and reducing ceroid lipofuscin accumulation in mouse hippocampus and somatosensory brain regions (Johnson et al., 2023). The auditory duration MMN paradigm provides a means to assess the functional impact of such therapies.

The observed alterations of auditory neurophysiological responses in *Cln3-/-* mice likely stem from sex-specific disease progression, since WT mice of both sexes showed little to no age-related changes in auditory duration MMN and AEPs within the age range (3-9 months) studied. The mechanisms underlying late-age compensation of auditory deficits in male *Cln3-/-* mice, but not in females, present an intriguing direction for future investigation. One potential avenue is to examine the relationship between ceroid lipofuscin accumulation — a pathological hallmark of CLN3 disease (Wright et al., 2020) — and auditory neurophysiological deficits. Because ceroid lipofuscin storage material contains a significant amount of the subunit C of mitochondrial ATP synthase (SCMAS) (Mitchison et al., 1999), SCMAS is widely used as a histopathological marker for CLN3 disease (Gomez-Giro et al., 2019). While previous research did not specifically examine auditory brain regions for SCMAS expression in *Cln3* mouse models, one recent study suggested that male *Cln3* mutant mice accumulate significantly more SCMAS in visual cortex, somatosensory cortex, thalamus, and hippocampus at postnatal day 70, compared to female *Cln3* mutant mice (Centa et al., 2023). The more pronounced auditory deficits in early-age male *Cln3-/-* mice may result from storage material accumulation exceeding their female counterparts. This highlights the importance of considering sex differences in the relationship between CLN3 disease progression related neuropathological phenotypes and neurophysiological impairments.

Exploring neuropathological markers can also provide potential insights for late-age compensation of auditory deficits in male *Cln3-/-* mice, but not in female *Cln3-/-* mice. However, sex differences in storage material accumulation remain underexamined at older ages in *Cln3* mouse models (McShane and Mole, 2022). Studies in the *Cln6* model of Batten disease revealed that male mutant mice accumulate more storage material in the somatosensory cortex and thalamus at 2 months of age, whereas female *Cln6* mutant mice surpass males in storage material by 6 months of age in these same regions (Poppens et al., 2019). A similar progression in *Cln3* mutant females could potentially explain the irreversible auditory dysfunctions observed at later ages in this study. However, previous studies also suggested that the accumulation of ceroid lipofuscin alone does not necessarily correlate with neural or behavioral deficits (Mitchison et al., 1999; Cotman et al., 2002). Further investigation into the neuropathology and pathophysiology in *Cln3-/-* mice along auditory processing pathways may help pinpoint specific regions and cell types involved in central auditory processing deficits and compensatory mechanisms. Importantly, identification of potential compensation processes in male *Cln3-/-* mice in the current study suggests a novel research direction to explore for therapeutic development.

This study bridges the gap between current pre-clinical and clinical research in CLN3 disease. Widely used assays in *Cln3* mouse models, such as invasive electrophysiological recording (Burkovetskaya et al., 2017) and behavioral assessments, ranging from maze tasks (Wendt et al., 2005) to open field (Centa et al., 2023) and rotarod tests (Osorio et al., 2009), are not readily translatable or comparable to human research. Importantly, our previous work identified auditory duration MMN deficits in individuals with CLN3 disease (Brima et al., 2024), and the current study demonstrates highly consistent abnormalities in *Cln3-/-* mice. This validates auditory duration MMN as a translational neurophysiological biomarker, enabling cross-species evaluation and comparison of sex- and age-specific progression in pathophysiological phenotypes. For instance, the drastic reductions of auditory duration MMN and AEPs in old female *Cln3-/-* mice has important clinical relevance, as female patients with CLN3 disease experience rapid progression towards severe symptoms, despite later onset compared to males (Munroe et al., 1997; Cialone et al., 2012; Nielsen and Ostergaard, 2013).

Auditory duration MMN may provide a more accurate means to evaluate treatment efficacy in CLN3 disease. For instance, an immunosuppressive agent, mycophenolate, showed efficacy in improving motor functions and attenuating neuroinflammation in *Cln3* mouse model (Seehafer et al., 2011). However, short-term administration of mycophenolate in patients with CLN3 disease showed no definite clinical outcomes (Augustine et al., 2019). Since auditory duration MMN paradigm is reliable for long-term assessment across species, it can be utilized for both forward and backward translation of pharmacological agents and verify both pre-clinical and clinical outcomes longitudinally.

Moreover, EEG recordings with the auditory duration MMN paradigm are appliable to other neurodevelopmental disorders. Previous work in Rett Syndrome (Brima et al., 2019), 22q11.2 deletion syndrome (Francisco et al., 2020b), and Cystinosis (Francisco et al., 2020a; Francisco et al., 2021) has identified MMN deficits in human participants. Expanding MMN studies to murine models of these disorders could facilitate longitudinal tracking of disease progression and exploration of sex and age differences. Validation of MMN as a translational neurophysiological biomarker across disorders may also aid in establishing common endophenotypes for various neurodevelopmental conditions.

A limitation of the current study is related to the spatial resolution of EEG. Given that EEG is a measure of synchronized electrical activity from large neuronal populations (Light et al., 2010), it is difficult to pinpoint the exact brain location underlying auditory duration MMN and AEP differences in WT and *Cln3-/-* mice. Since MMN is a higher-order brain response generated from interactions between auditory thalamocortical circuits and frontal-temporal brain regions (Lakatos et al., 2020), further investigations using complementary approaches are needed to elucidate the neural mechanisms driving these deficits.

In conclusion, WT mice of both sexes maintained normal central auditory processing over time, whereas male *Cln3-/-* mice showed restoration of early deficits and female *Cln3-/-* mice exhibited a progressive decline. These findings provide a foundation to characterize sex- and disease progression-specific sensory processing deficits, uncover underlying neural mechanisms, and utilize auditory duration MMN as a cross-species neurophysiological biomarker.

## Acknowledgment

This research is funded by the National Institutes of Health (P50 HD103536-7954 to KHW and JJF). We thank members of Wang Lab and Frederick J. and Marion A. Schindler Cognitive Neurophysiology Lab for critical discussions.

