## Supplementary figures for "Sex-specific and age-related progression of auditory neurophysiological deficits in the *Cln3* mouse model of Batten disease"

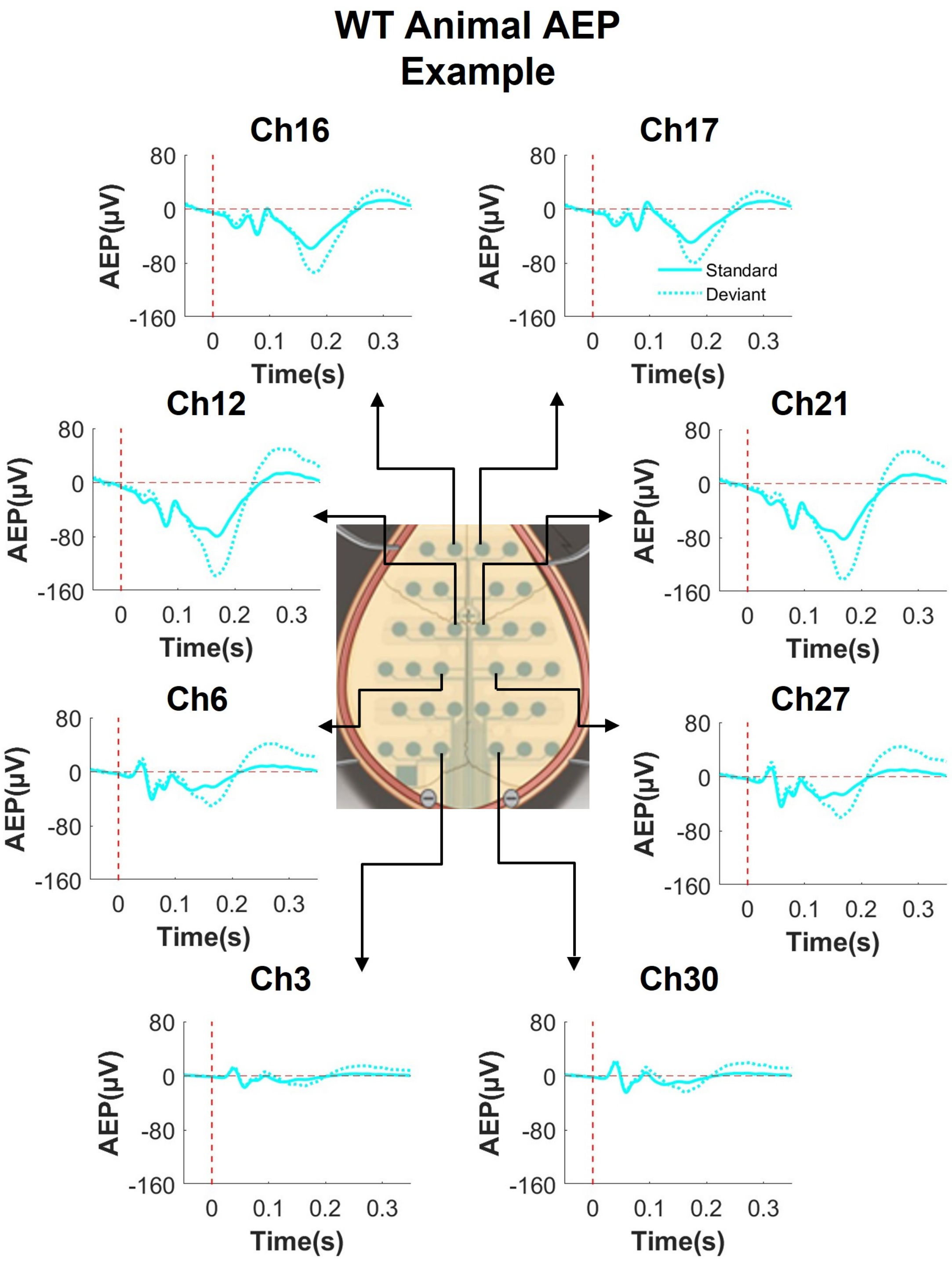
WT Animal AEP
Example
Ch16
Ch17
Ch12
Ch21
Ch6
Ch27
Ch3
Ch30
AEP(µV)
Time(s)
Standard
Deviant
80
0
-80
-160
0
0.1
0.2
0.3

**Extended Data Figure 1-1. EEG mice showed robust responses to both standard and deviant stimuli in spatial distribution.** Standard and deviant AEP waveforms from one representative WT male mouse. Ch stands for channel number. The vertical dash red line indicates the onset of stimuli, and the horizontal dash red line indicates 0. Averaged AEP to standard tones is presented with solid cyan line and averaged AEP to deviant tones is presented with dotted cyan line. The example mouse showed clear AEPs to both standard and deviant tones from anterior to posterior region over the scalp, with MMN seen maximal at the center channels (i.e., Ch12 and Ch21). AEPs have left-right hemisphere symmetry. AEPs at most posterior channels (i.e., Ch3 and Ch30) are smaller due to referencing.

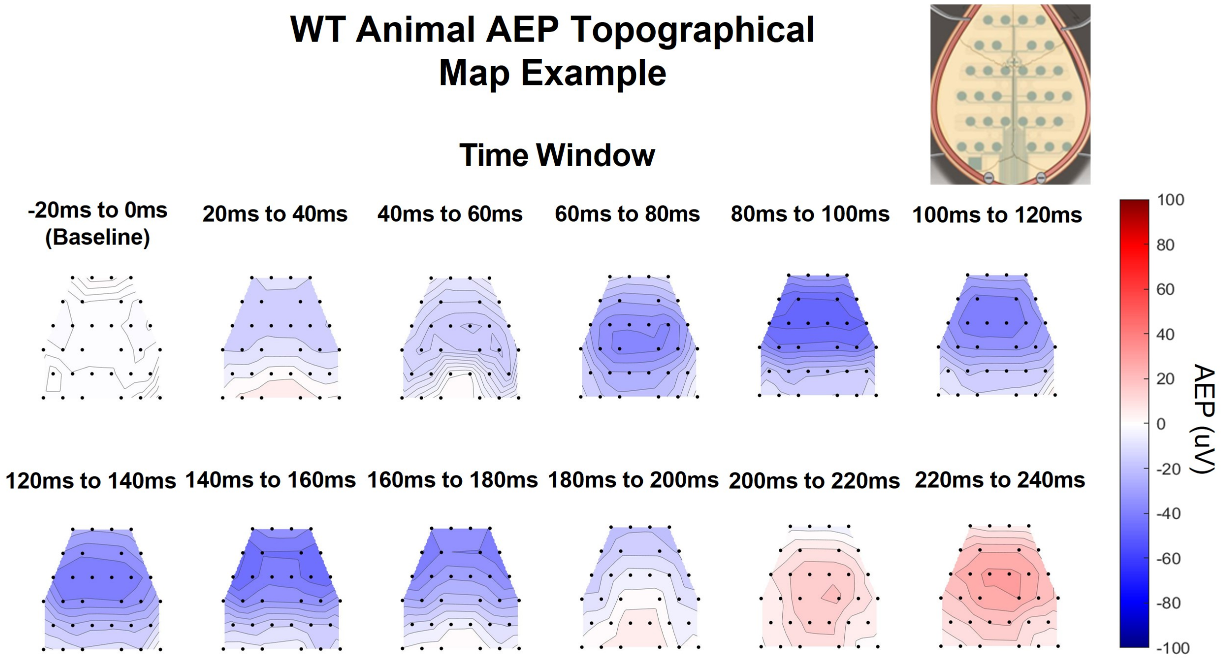


**Extended Data Figure 1-2. EEG mice showed clear propagation of AEPs in spatial distribution.** Topographical maps of standard AEP in a 20ms time window from one representative WT male mouse. Black dots indicate the position of EEG electrodes. The color bar shows the positivity (red) and negativity (blue) of the AEP response. AEPs in WT animals propagate from temporal regions near the Auditory Cortex (AC) to central parietal regions, and subsequently to frontal regions.

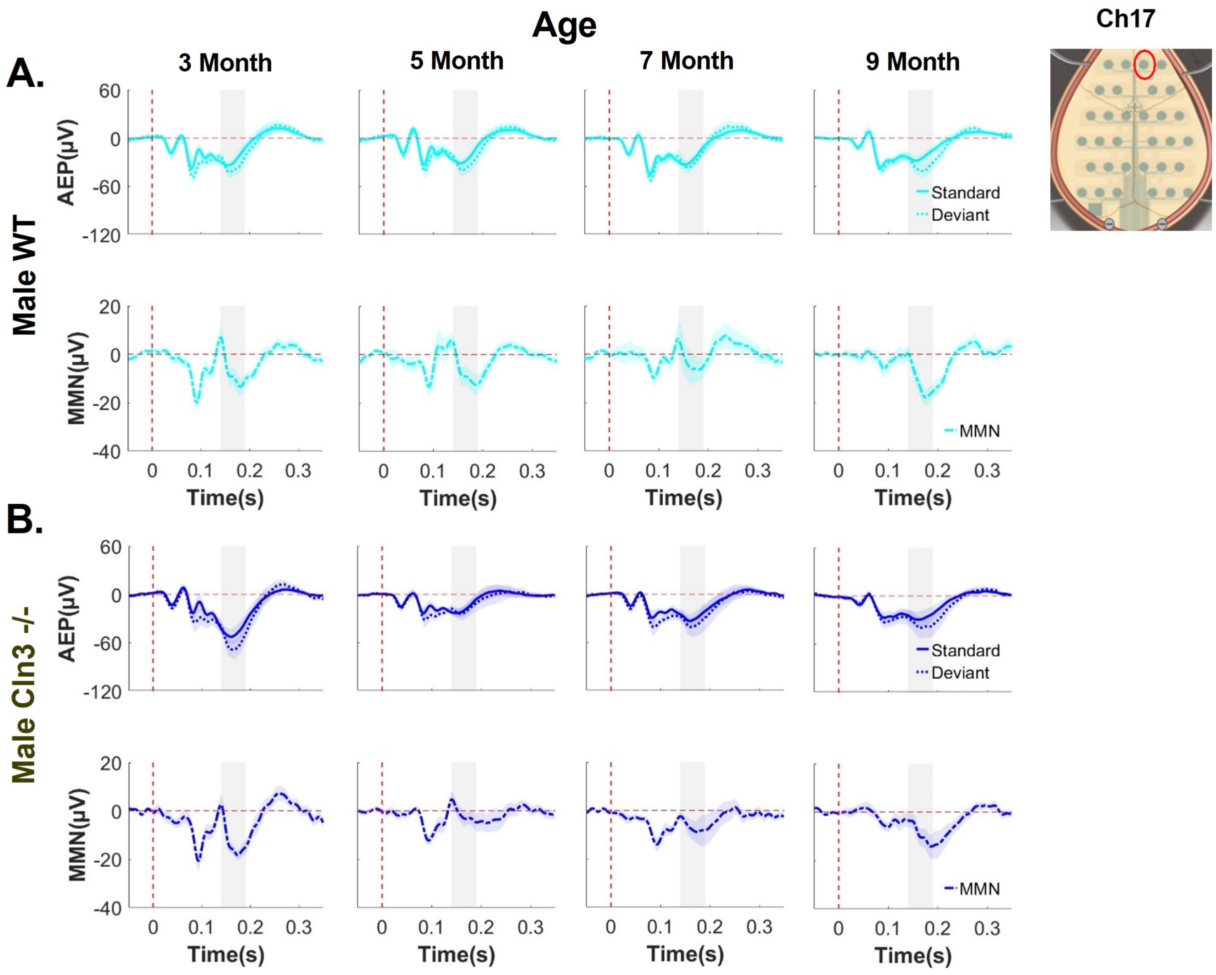


**Extended Data Figure 3-1. Male WT mice showed robust responses, while male *Cln3-/-* mice showed disease progression-specific deficit of auditory duration MMN across different electrodes.** Group-averaged AEP and auditory duration MMN waveforms for male WT and *Cln3-/-* mice at electrode 17 (most anterior). Vertical dash red line indicates onset of stimuli and horizontal dash red line indicates 0. Standard AEP in solid line, deviant AEP in dotted line, and MMN in solid- dotted line. Male WT mice (n = 11 for 3-, 7- and 9-month, n = 14 for 5 month) in light blue, male *Cln3-/-* mice (n = 8 for 3-, 7- and 9-month, n = 11 for 5 month) in dark blue.

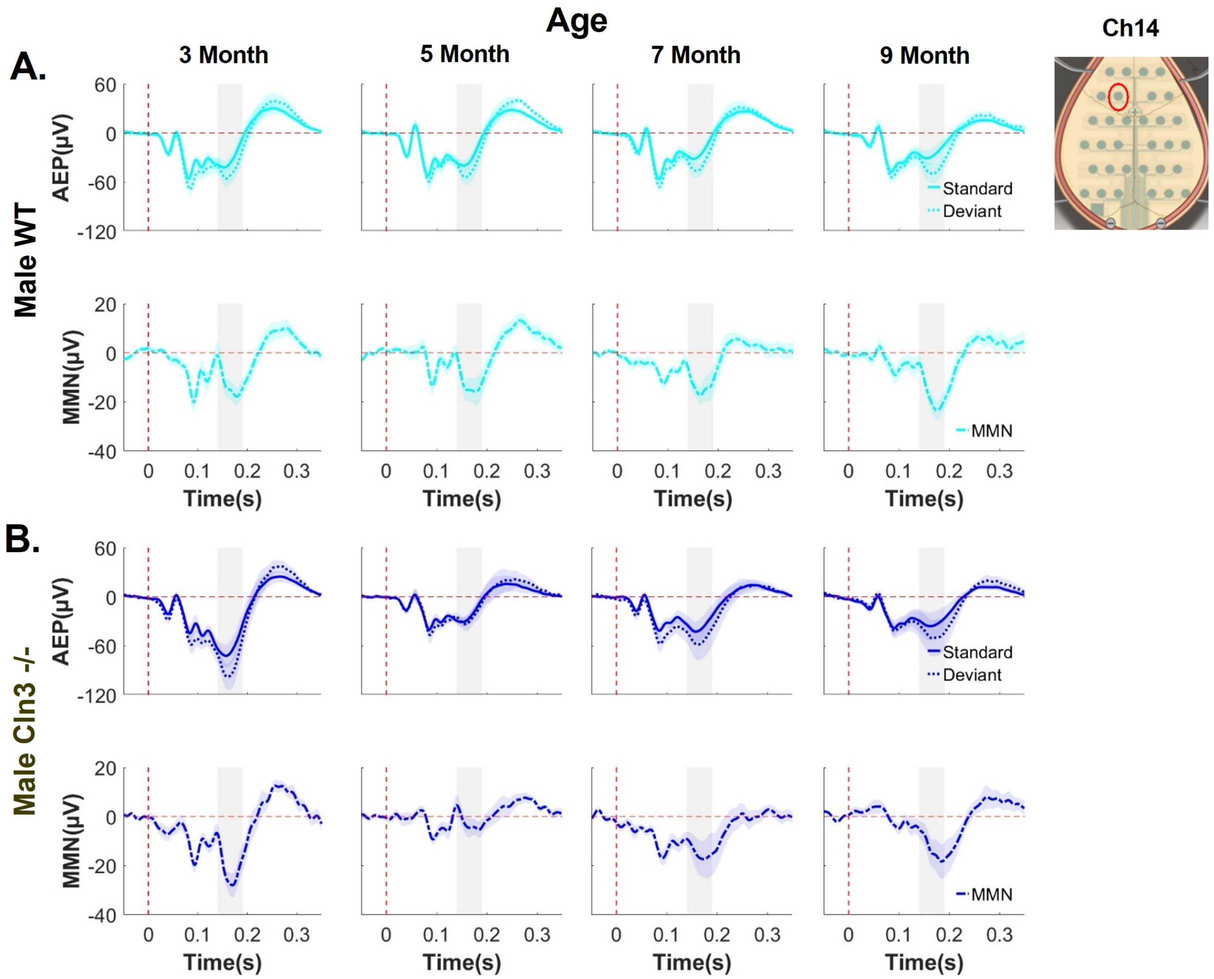


**Extended Data Figure 3-2. Male WT mice showed robust response, while male *Cln3-/-* mice showed disease progression-specific deficit of auditory duration MMN across different electrodes.** Group-averaged AEP and auditory duration MMN waveforms for male WT and *Cln3-/-* mice at electrode 14 (anterior). The vertical dash red line indicates the onset of stimuli, and the horizontal dash red line indicates 0. Standard AEP in a solid line, deviant AEP in a dotted line, and MMN in a solid- dotted line. Male WT mice (n = 11 for 3, 7 and 9 months, n = 14 for 5 months) in light blue, male *Cln3-/-* mice (n = 8 for 3, 7 and 9 months, n = 11 for 5 months) in dark blue.

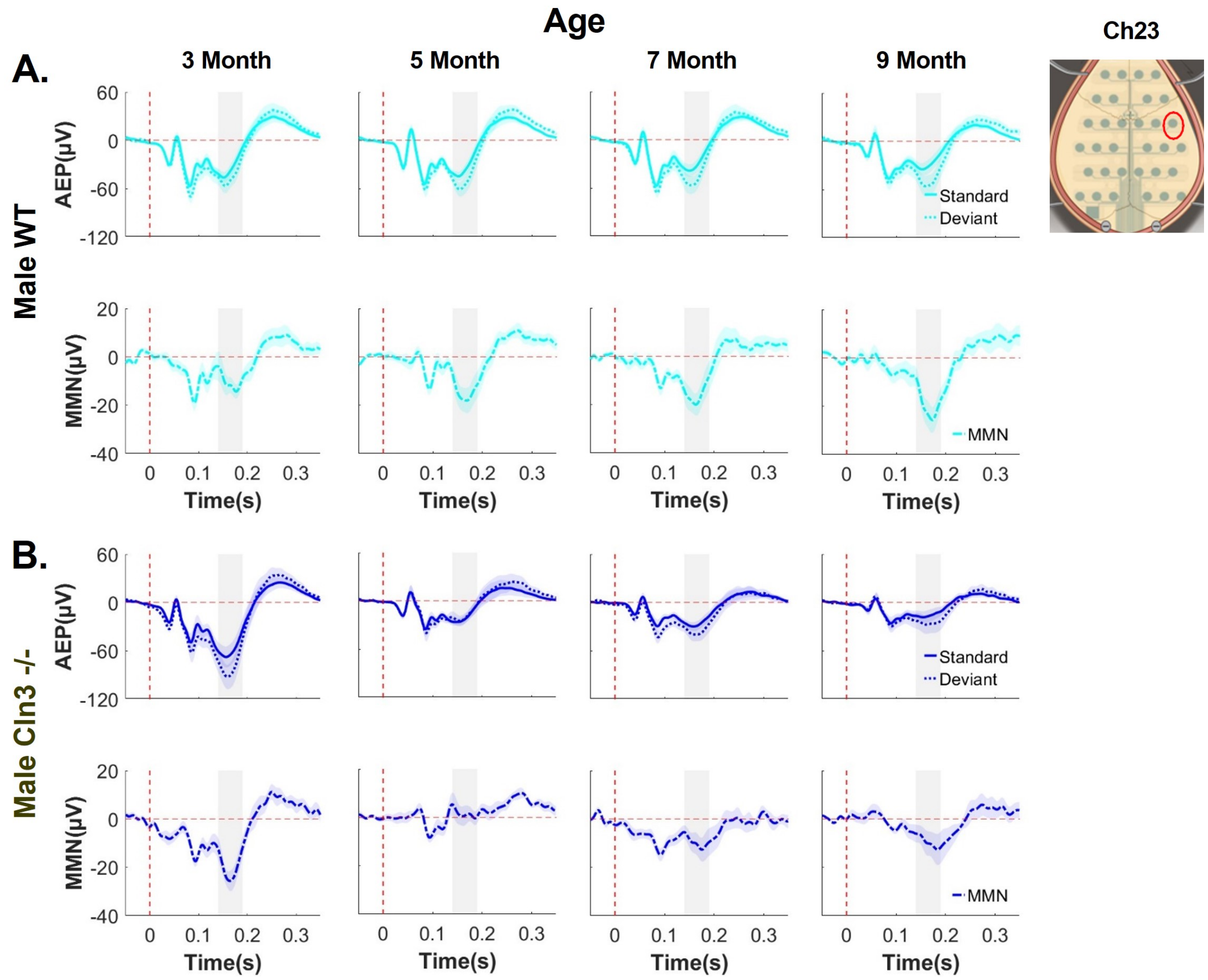


**Extended Data Figure 3-3. Male WT mice showed robust responses, while male *Cln3-/-* mice showed disease progression-specific deficit of auditory duration MMN across different electrodes.** **G**roup-averaged AEP and auditory duration MMN waveforms for male WT and *Cln3-/-* mice at electrode 23 (central). The vertical dash red line indicates the onset of stimuli, and the horizontal dash red line indicates 0. Standard AEP in a solid line, deviant AEP in dotted line, and MMN in a solid- dotted line. Male WT mice (n = 11 for 3, 7, and 9 months, n = 14 for 5 months) in light blue, male *Cln3-/-* mice (n = 8 for 3, 7 and 9 months, n = 11 for 5 month) in dark blue.

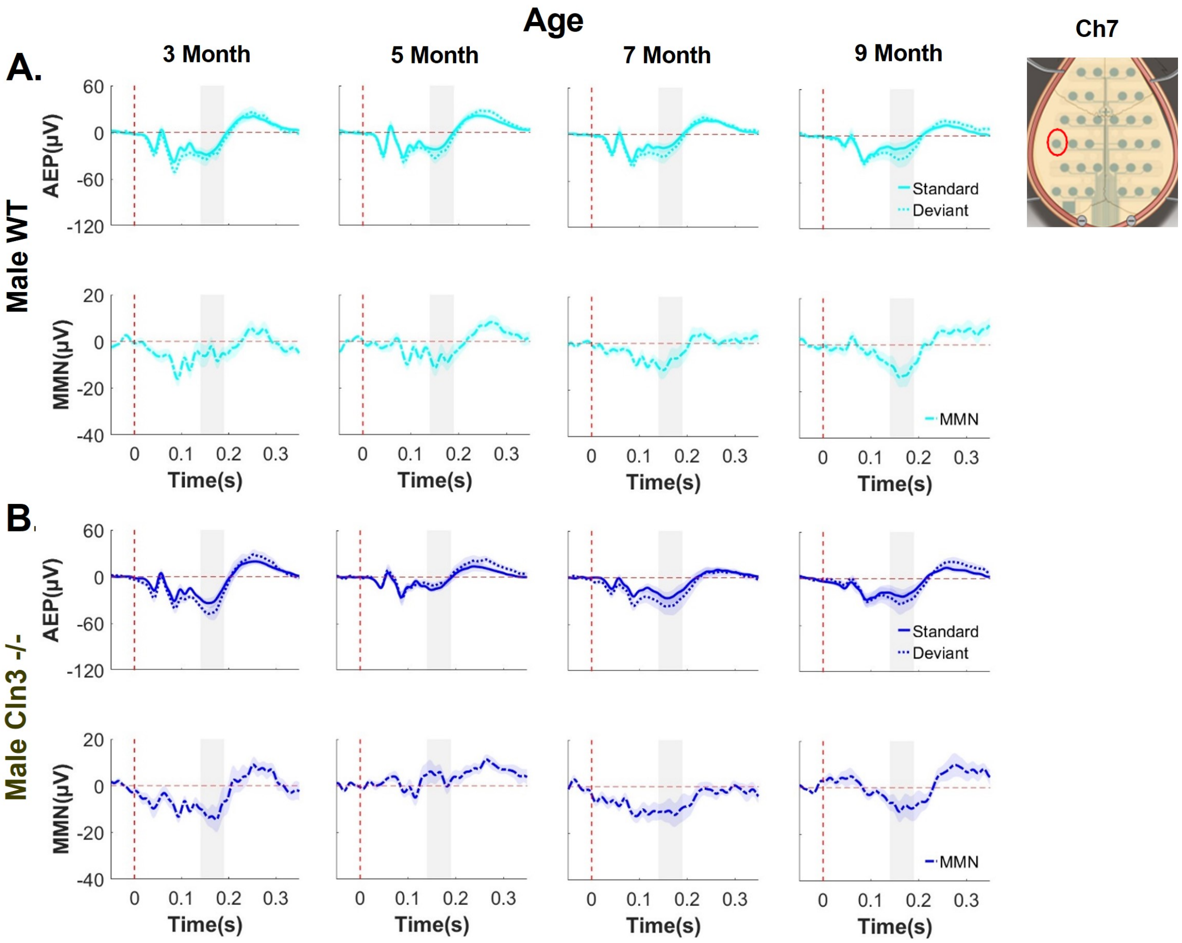


**Extended Data Figure 3-4. Male WT mice showed robust responses, while male *Cln3-/-* mice showed disease progression-specific deficit of auditory duration MMN across different electrodes.** Group-averaged AEP and auditory duration MMN waveforms for male WT and *Cln3-/-* mice at electrode 7 (central). The vertical dash red line indicates the onset of stimuli, and the horizontal dash red line indicates 0. Standard AEP in a solid line, deviant AEP in a dotted line, and MMN in dotted line. Male WT mice (n = 11 for 3, 7, and 9 months, n = 14 for 5 months) in light blue, male *Cln3-/-* mice (n = 8 for 3, 7, and 9 months, n = 11 for 5 month) in dark blue.

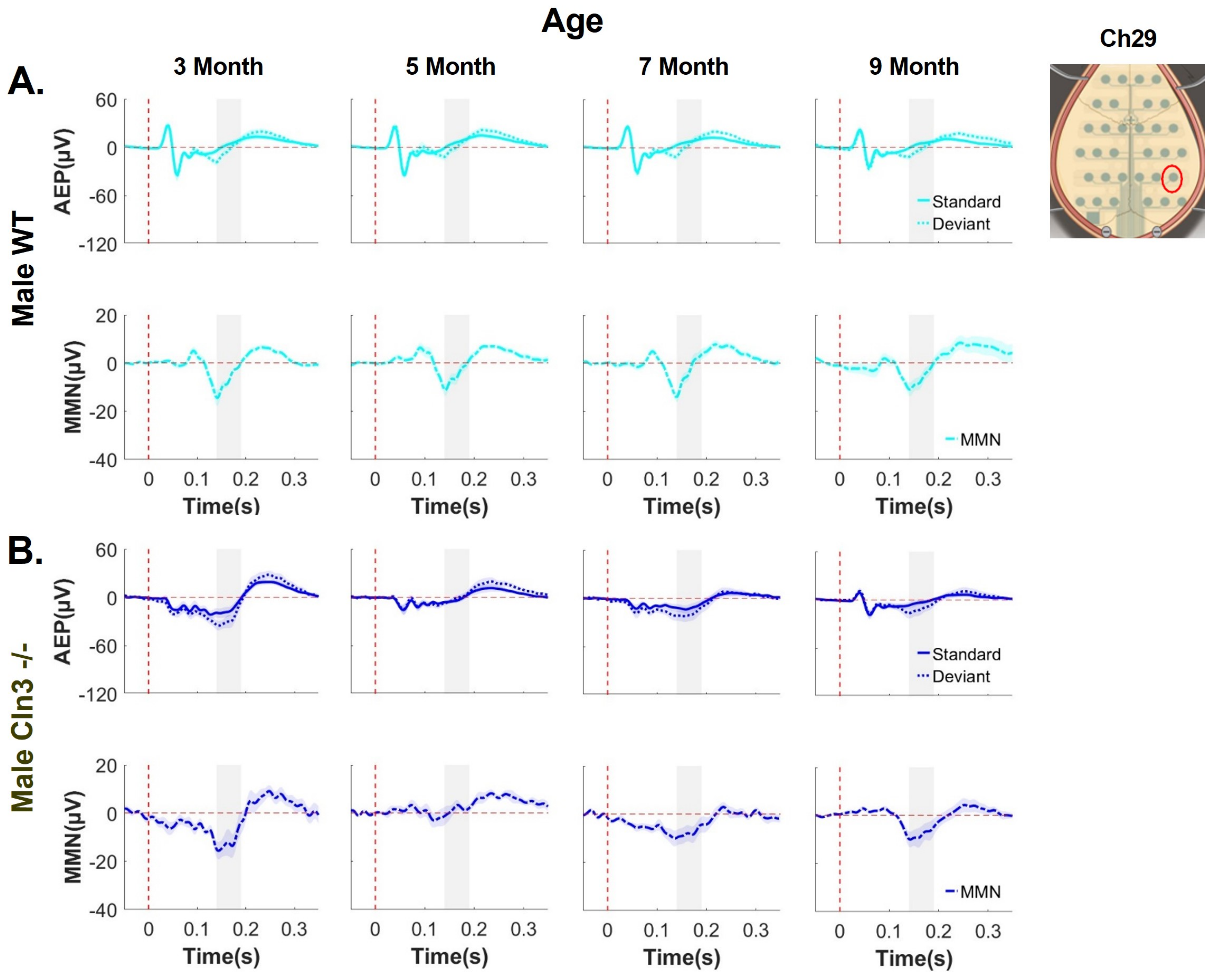


**Extended Data Figure 3-5. Male WT mice showed robust responses, while male *Cln3-/-* mice showed disease progression-specific deficit of auditory duration MMN across different electrodes.** Group-averaged AEP and auditory duration MMN waveforms for male WT and *Cln3-/-* mice at electrode 29 (posterior). The vertical dash red line indicates the onset of stimuli, and the horizontal dash red line indicates 0. Standard AEP in a solid line, deviant AEP in a dotted line, and MMN in a solid-dotted line. Male WT mice (n = 11 for 3, 7, and 9 months, n = 14 for 5 months) in light blue, male *Cln3-/-* mice (n = 8 for 3, 7, and 9 months, n = 11 for 5 month) in dark blue.

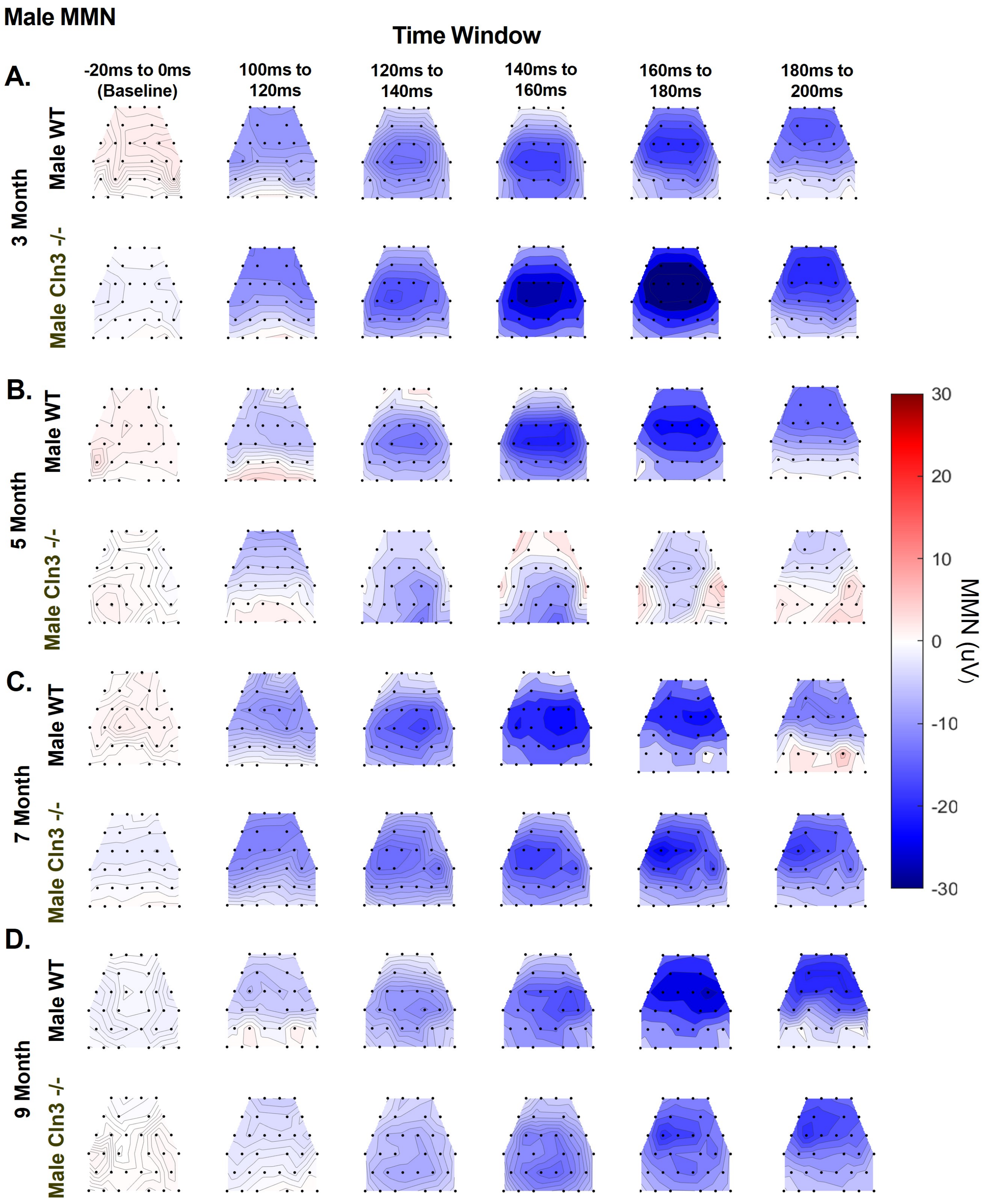
Male MMN
Time Window
A.
-20ms to 0ms (Baseline)
100ms to 120ms
120ms to 140ms
140ms to 160ms
160ms to 180ms
180ms to 200ms
3 Month
Male WT
Male Cln3 -/-
B.
5 Month
Male WT
Male Cln3 -/-
C.
7 Month
Male WT
Male Cln3 -/-
D.
9 Month
Male WT
Male Cln3 -/-
30
20
10
0
-10
-20
-30
MMN (uV)

**Extended Data Figure 3-6. Male *Cln3-/-* mice showed initial greater response, then decline and later recovery of auditory duration MMN in spatial distribution.** Averaged topographical maps of auditory duration MMN in a 20ms time window from male WT (n = 11 for 3, 7, and 9 months, n = 14 for 5 months) and male *Cln3-/-* mice (n = 8 for 3, 7 and 9 months, n = 11 for 5 months). ***A*** MMN topographical maps for 3-month-old mice. ***B,*** MMN topographical maps for 5-month-old mice. ***C,*** MMN topographical maps for 7-month-old mice. ***D,*** MMN topographical maps for 9 months old mice. Male *Cln3-/-* mice showed initial greater response, then decline and later recovery of auditory duration MMN in spatial distribution.

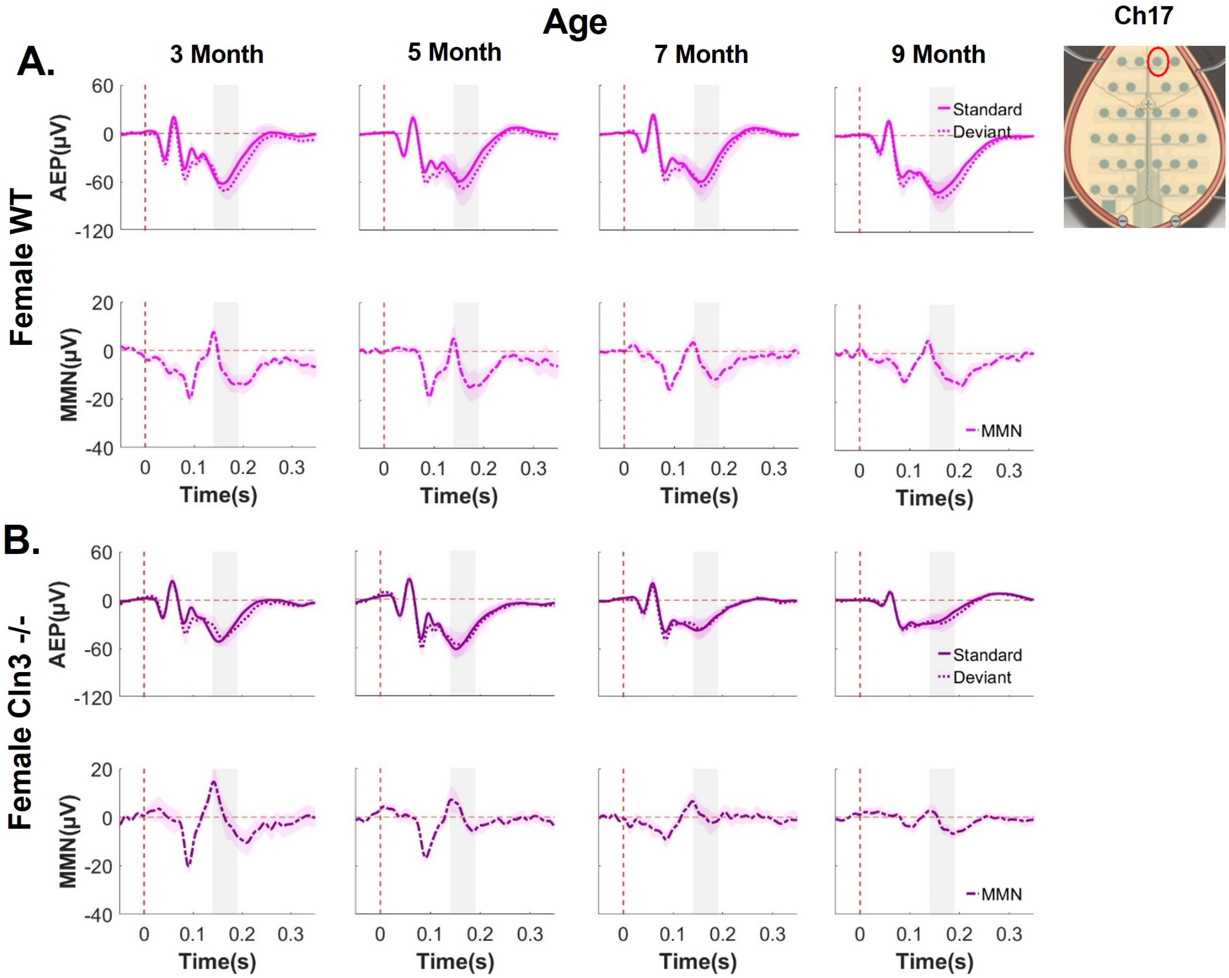


**Extended Data Figure 4-1. Female WT mice showed robust responses, while female *Cln3-/-* mice showed a progressive decline in auditory duration MMN across different electrodes.** Group-averaged AEP and auditory duration MMN waveforms for female WT and *Cln3-/-* mice at electrode 17 (most anterior). The vertical dash red line indicates the onset of stimuli, and the horizontal dash red line indicates 0. Standard AEP in a solid line, deviant AEP in a dotted line, and MMN in solid- dotted line. Female WT mice (n = 6 for all ages) in light pink and female Cln3-/- mice (n = 7 for all ages) in dark pink.

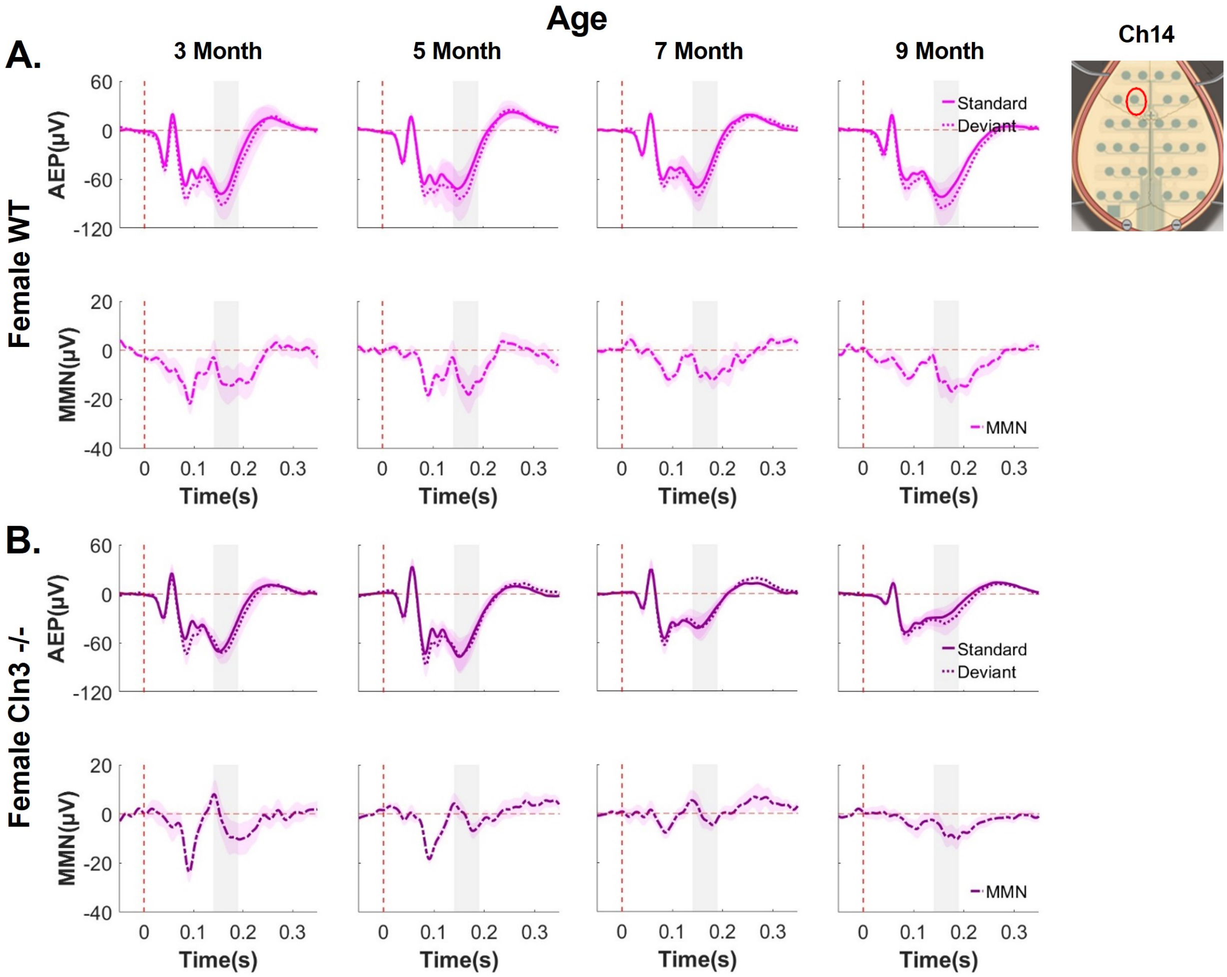


**Extended Data Figure 4-2. Female WT mice showed robust responses, while female *Cln3-/-* mice showed a progressive decline in auditory duration MMN across different electrodes.** Group-averaged AEP and auditory duration MMN waveforms for female WT and *Cln3-/-* mice at electrode 14 (anterior). The vertical dash red line indicates the onset of stimuli, and the horizontal dash red line indicates 0. Standard AEP in a solid line, deviant AEP in a dotted line, and MMN in a solid-dotted line. Female WT mice (n = 6 for all ages) in light pink and female Cln3-/- mice (n = 7 for all ages) in dark pink.

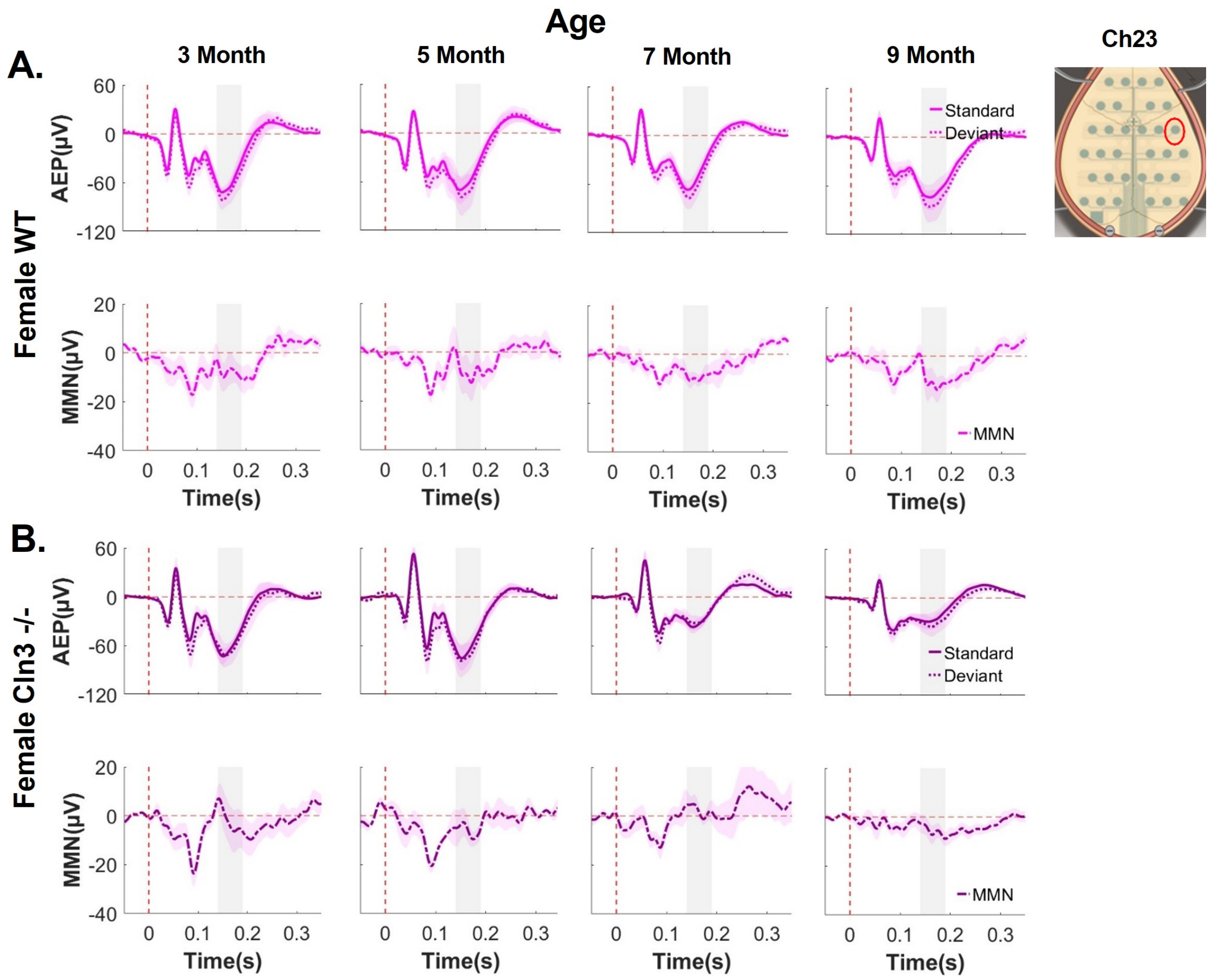


**Extended Data Figure 4-3. Female WT mice showed robust responses, while female *Cln3-/-* mice showed a progressive decline in auditory duration MMN across different electrodes.** Group-averaged AEP and auditory duration MMN waveforms for female WT and *Cln3-/-* mice at electrode 23 (central). The vertical dash red line indicates the onset of stimuli, and the horizontal dash red line indicates 0. Standard AEP in a solid line, deviant AEP in a dotted line, and MMN in a solid-dotted line. Female WT mice (n = 6 for all ages) in light pink and female Cln3-/- mice (n = 7 for all ages) in dark pink.

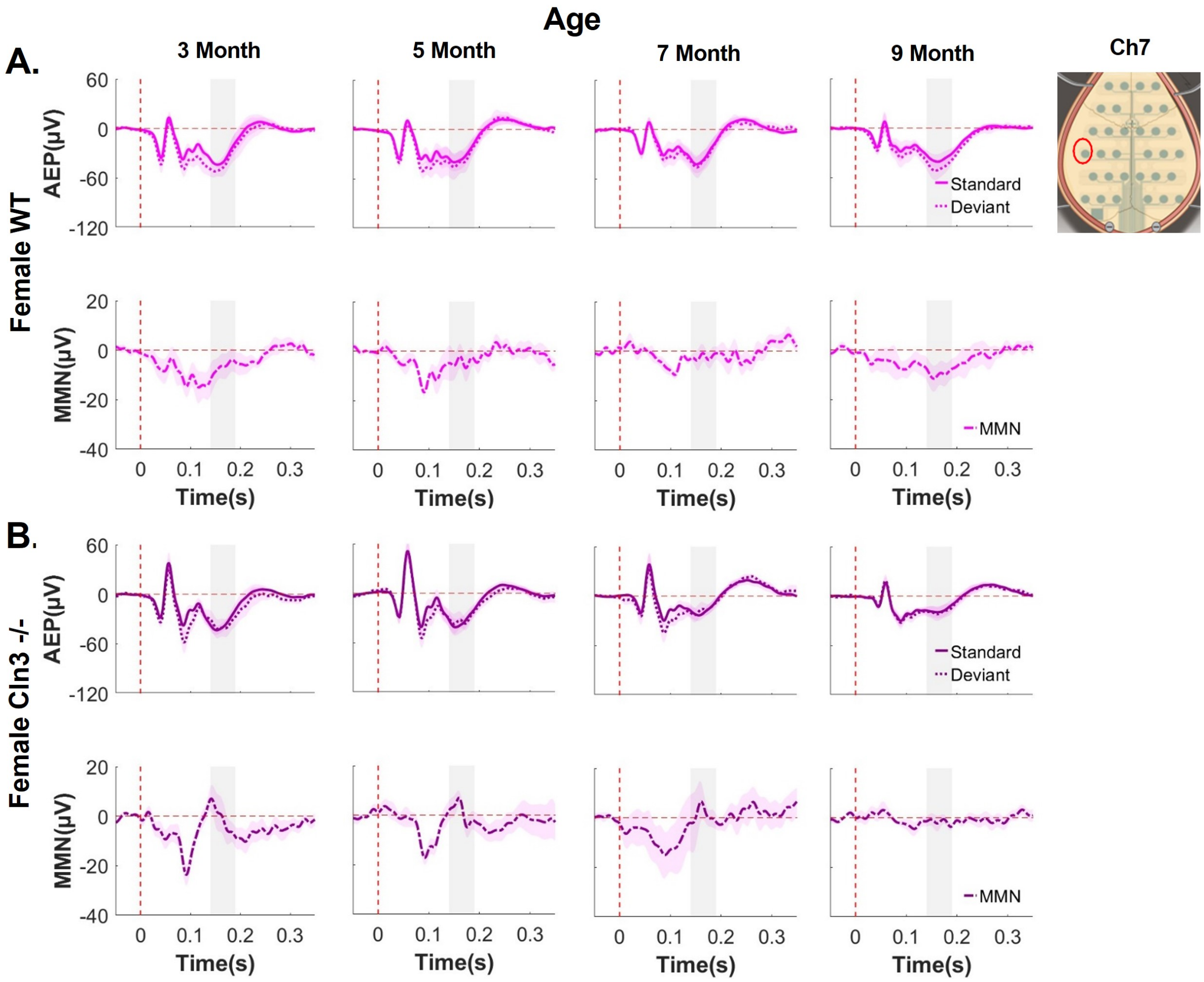


**Extended Data Figure 4-4. Female WT mice showed robust responses, while female *Cln3-/-* mice showed a progressive decline in auditory duration MMN across different electrodes.** Group-averaged AEP and auditory duration MMN waveforms for female WT and *Cln3-/-* mice at electrode 7 (central). The vertical dash red line indicates the onset of stimuli, and the horizontal dash red line indicates 0. Standard AEP in a solid line, deviant AEP in a dotted line, and MMN in a solid-dotted line. Female WT mice (n = 6 for all ages) in light pink and female Cln3-/- mice (n = 7 for all ages) in dark pink.

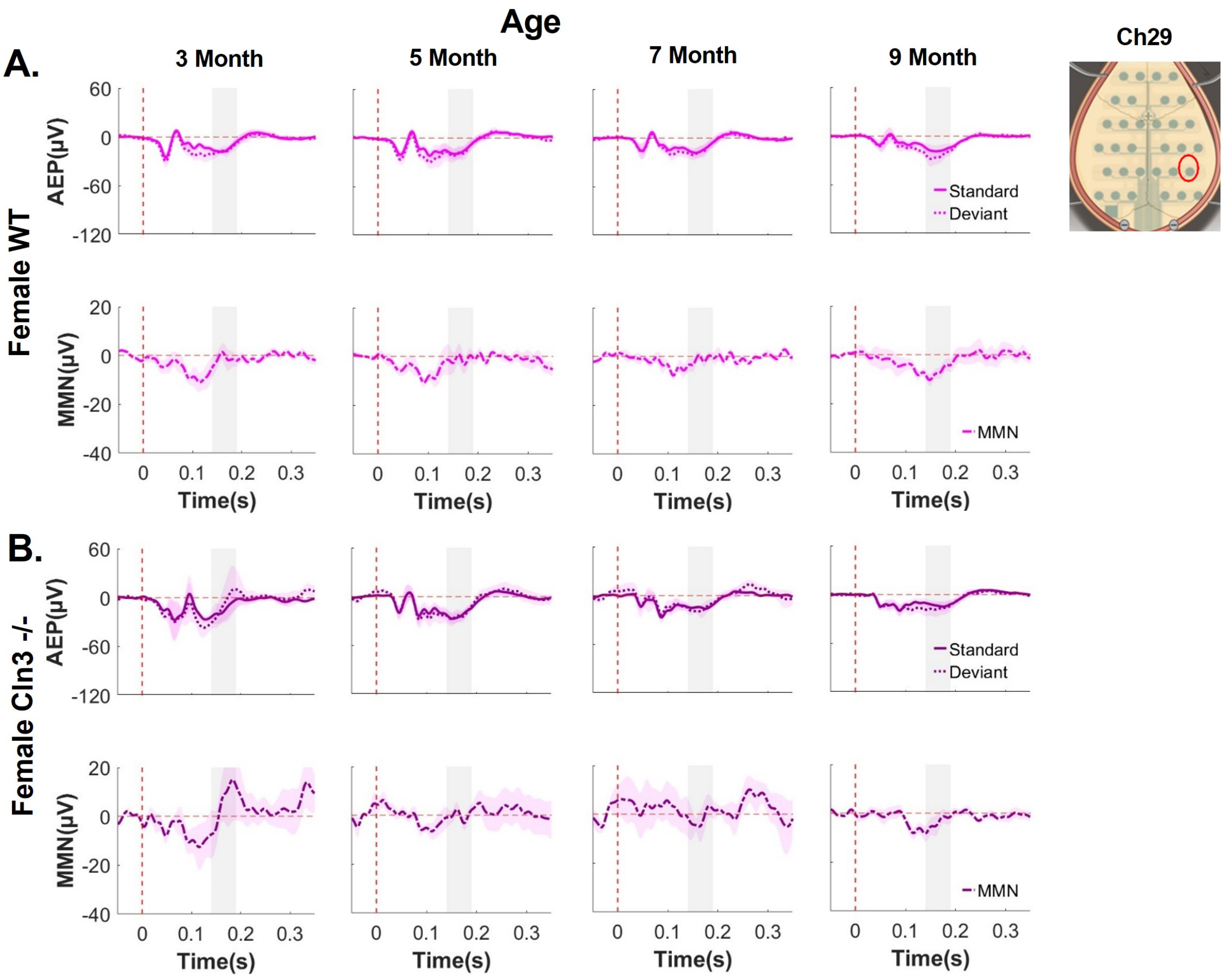


**Extended Data Figure 4-5. Female WT mice showed robust responses, while female *Cln3-/-* mice showed a progressive decline in auditory duration MMN across different electrodes.** Group-averaged AEP and auditory duration MMN waveforms for female WT and *Cln3-/-* mice at electrode 29 (posterior). The vertical dash red line indicates the onset of stimuli, and the horizontal dash red line indicates 0. Standard AEP in a solid line, deviant AEP in a dotted line, and MMN in a solid-dotted line. Female WT mice (n = 6 for all ages) in light pink and female Cln3-/- mice (n = 7 for all ages) in dark pink.

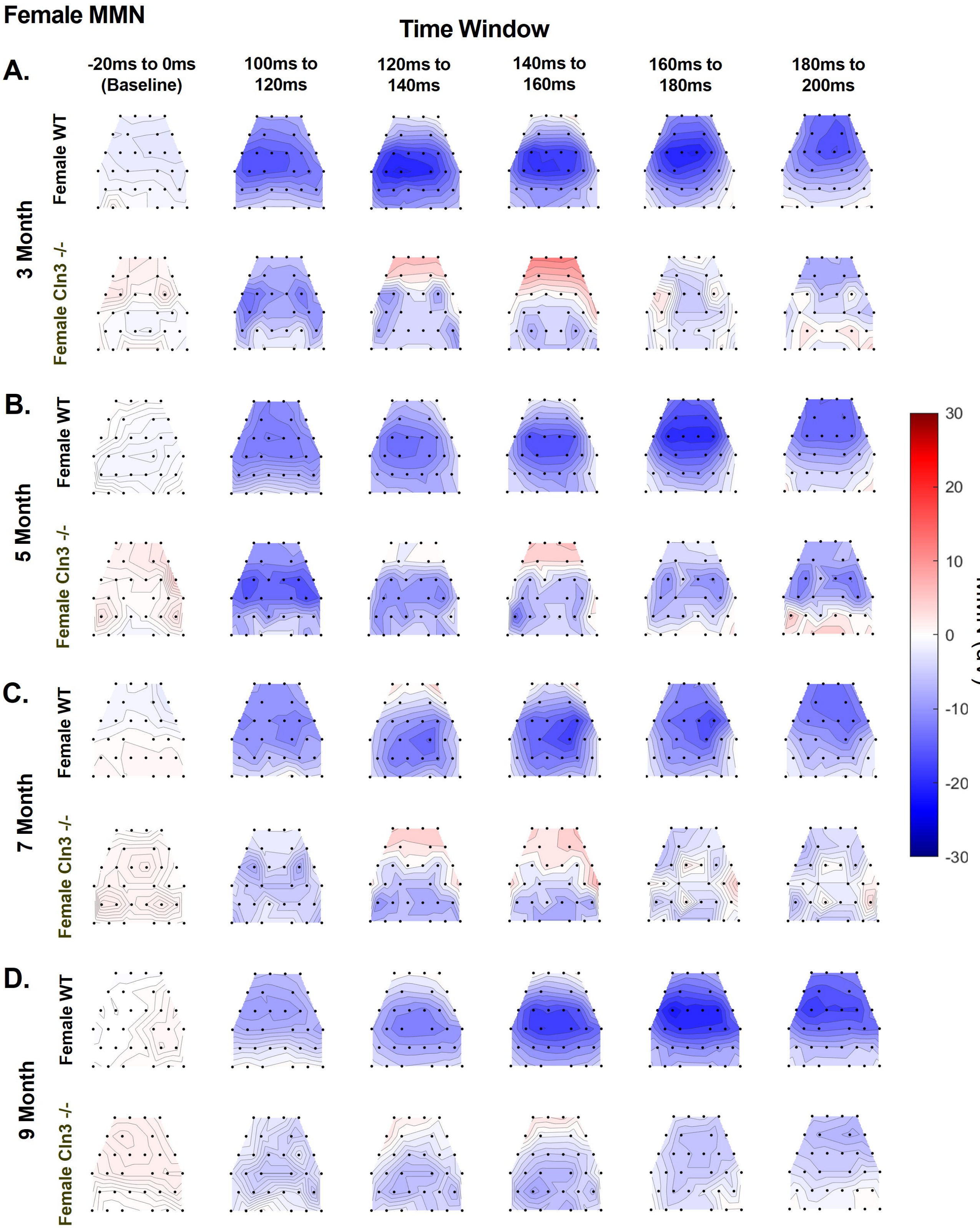

Female MMN
Time Window
A.
B.
C.
D.
-20ms to 0ms (Baseline)
100ms to 120ms
120ms to 140ms
140ms to 160ms
160ms to 180ms
180ms to 200ms
3 Month
5 Month
7 Month
9 Month
Female WT
Female Cln3 -/-
30
20
10
0
-10
-20
-30
MMN (uV)

**Extended Data Figure 4-6. Female *Cln3-/-* mice showed consistent auditory duration MMN in spatial distribution.** Averaged topographical maps of auditory duration MMN in a 20ms time window from female WT (n = 6 for all ages) and female *Cln3-/-* mice (n = 7 for all ages). ***A,*** MMN topographical maps for 3-month-old mice. ***B,*** MMN topographical maps for 5-month-old mice. ***C,*** MMN topographical maps for 7-month-old mice. ***D,*** MMN topographical maps for 9-month-old mice. Female *Cln3-/-* mice showed consistent auditory duration MMN in spatial distribution.

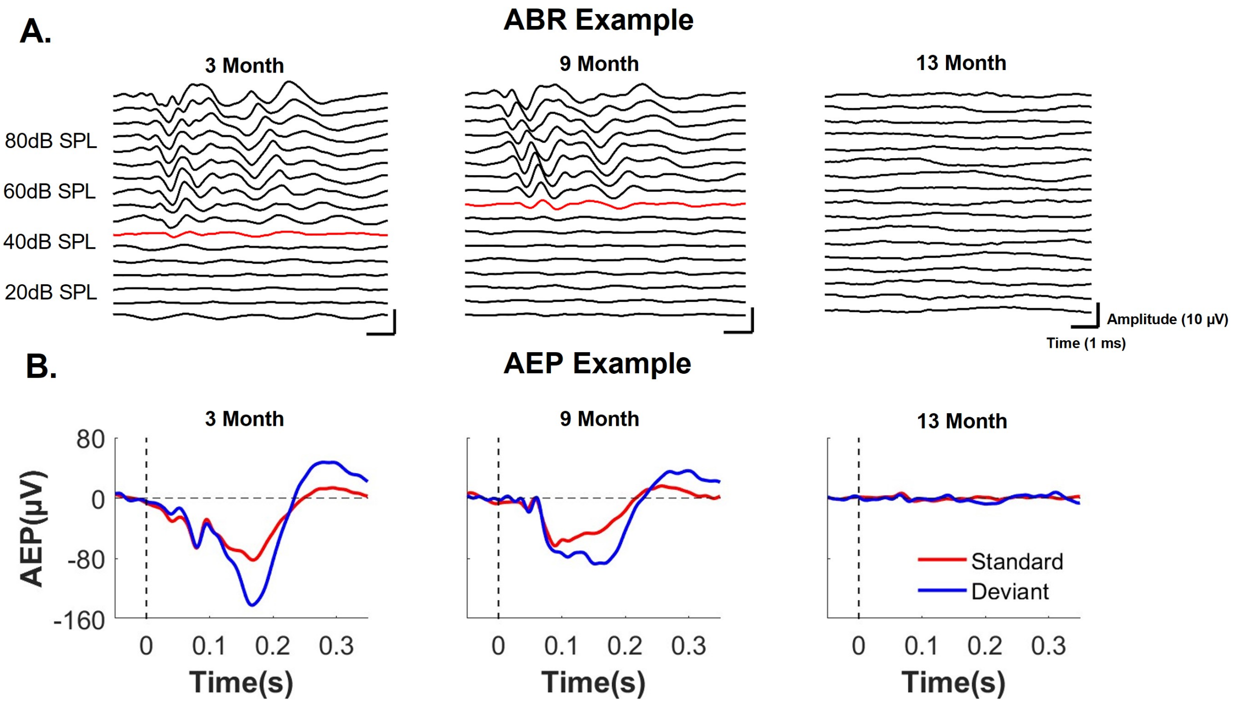


**Extended Data Figure 5-1. Animals with complete hearing loss had no identifiable ABR and AEP waveforms.** ***A,*** ABR waveforms from one 3-month-old, one 9-month-old, and one 13-month-old animal. There was no identifiable ABR waveform from the 13-month-old animal, suggesting a hearing threshold above 90dB SPL. ***B,*** group-averaged AEP waveforms for one 3-month-old, one 9-month-old, and one 13-month-old animal at Ch21. The vertical dash red line indicates the onset of stimuli, and the horizontal dash red line indicates 0. Standard AEP is in red and deviant AEP is in blue. Animals at younger ages showed clear AEPs with robust auditory duration MMN. There were no identifiable AEP waveforms for the 13-month-old animal. Complete hearing loss leads to the diminishment of ABRs and AEPs.

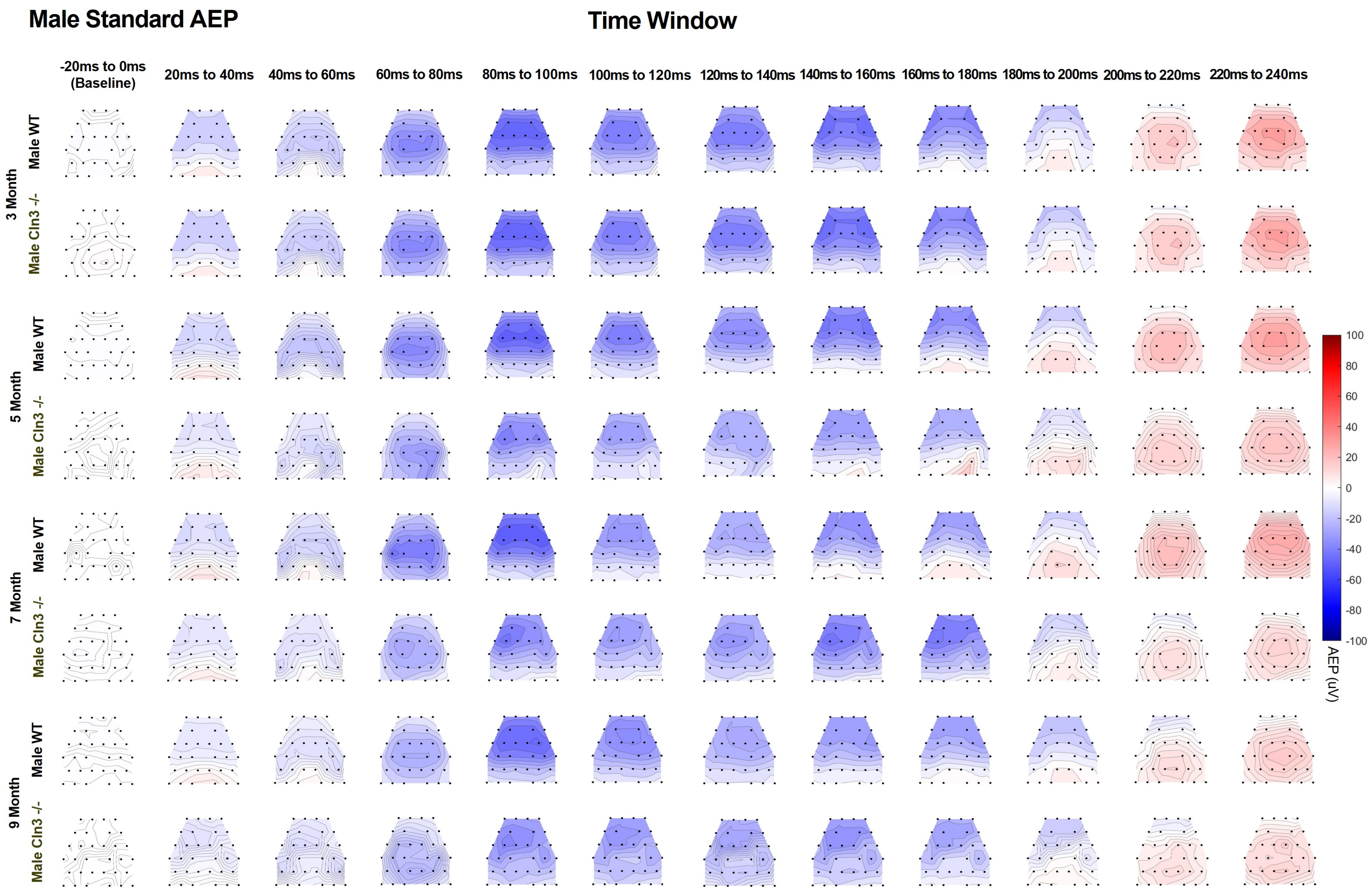

Male Standard AEP
Time Window
-20ms to 0ms (Baseline)
20ms to 40ms
40ms to 60ms
60ms to 80ms
80ms to 100ms
100ms to 120ms
120ms to 140ms
140ms to 160ms
160ms to 180ms
180ms to 200ms
200ms to 220ms
220ms to 240ms
3 Month
5 Month
7 Month
9 Month
Male WT
Male Cln3 -/-
100
80
60
40
20
0
-20
-40
-60
-80
-100
AEP (uV)

**Extended Data Figure 6-1. Male *Cln3-/-* mice showed initial greater response then decline and later recovery of AEPs in spatial distribution.** Averaged topographical maps of standard AEPs in a 20ms time window from male WT (n = 11 for 3, 7, and 9 months, n = 14 for 5 months) and male *Cln3-/-* mice (n = 8 for 3, 7 and 9 months, n = 11 for 5 months). Male *Cln3-/-* mice showed an initial greater response, then decline and later recovery of standard AEPs in spatial distribution. Blue shows negativity and red shows positivity for AEP.

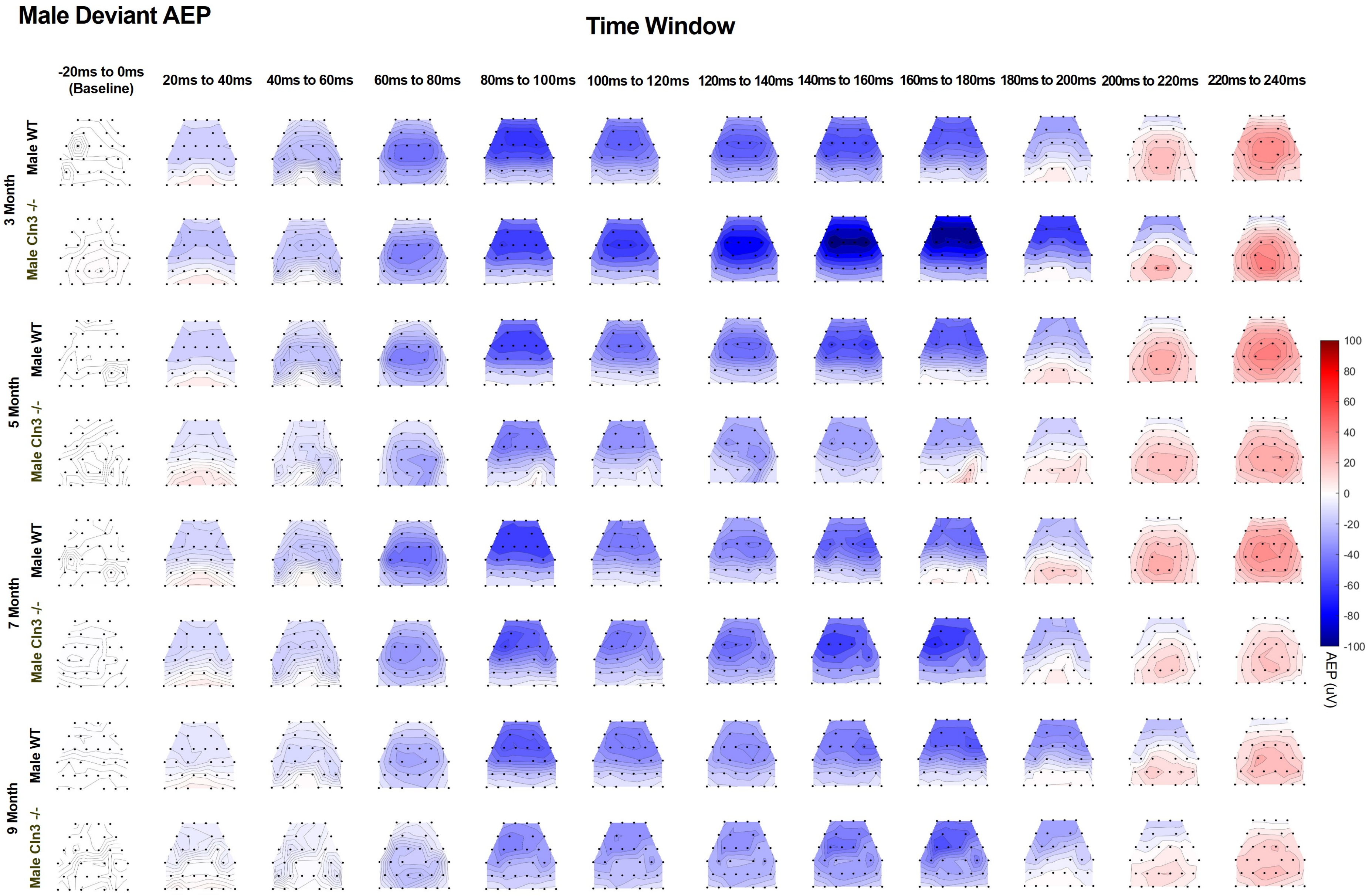
Male Deviant AEP
Time Window
-20ms to 0ms (Baseline)
20ms to 40ms
40ms to 60ms
60ms to 80ms
80ms to 100ms
100ms to 120ms
120ms to 140ms
140ms to 160ms
160ms to 180ms
180ms to 200ms
200ms to 220ms
220ms to 240ms
3 Month
Male WT
Male Cln3 -/-
5 Month
Male WT
Male Cln3 -/-
7 Month
Male WT
Male Cln3 -/-
9 Month
Male WT
Male Cln3 -/-
100
80
60
40
20
0
-20
-40
-60
-80
-100
AEP (uV)

**Extended Data Figure 6-2. Male *Cln3-/-* mice showed initial greater response then decline and later recovery of AEPs in spatial distribution.** Averaged topographical maps of deviant AEPs in a 20ms time window from male WT (n = 11 for 3, 7, and 9 months, n = 14 for 5 months) and male *Cln3-/-* mice (n = 8 for 3, 7, and 9 months, n = 11 for 5 months). Male *Cln3-/-* mice showed initial greater response, then decline and later recovery of deviant AEPs in spatial distribution. Blue shows negativity and red shows positivity for AEP.

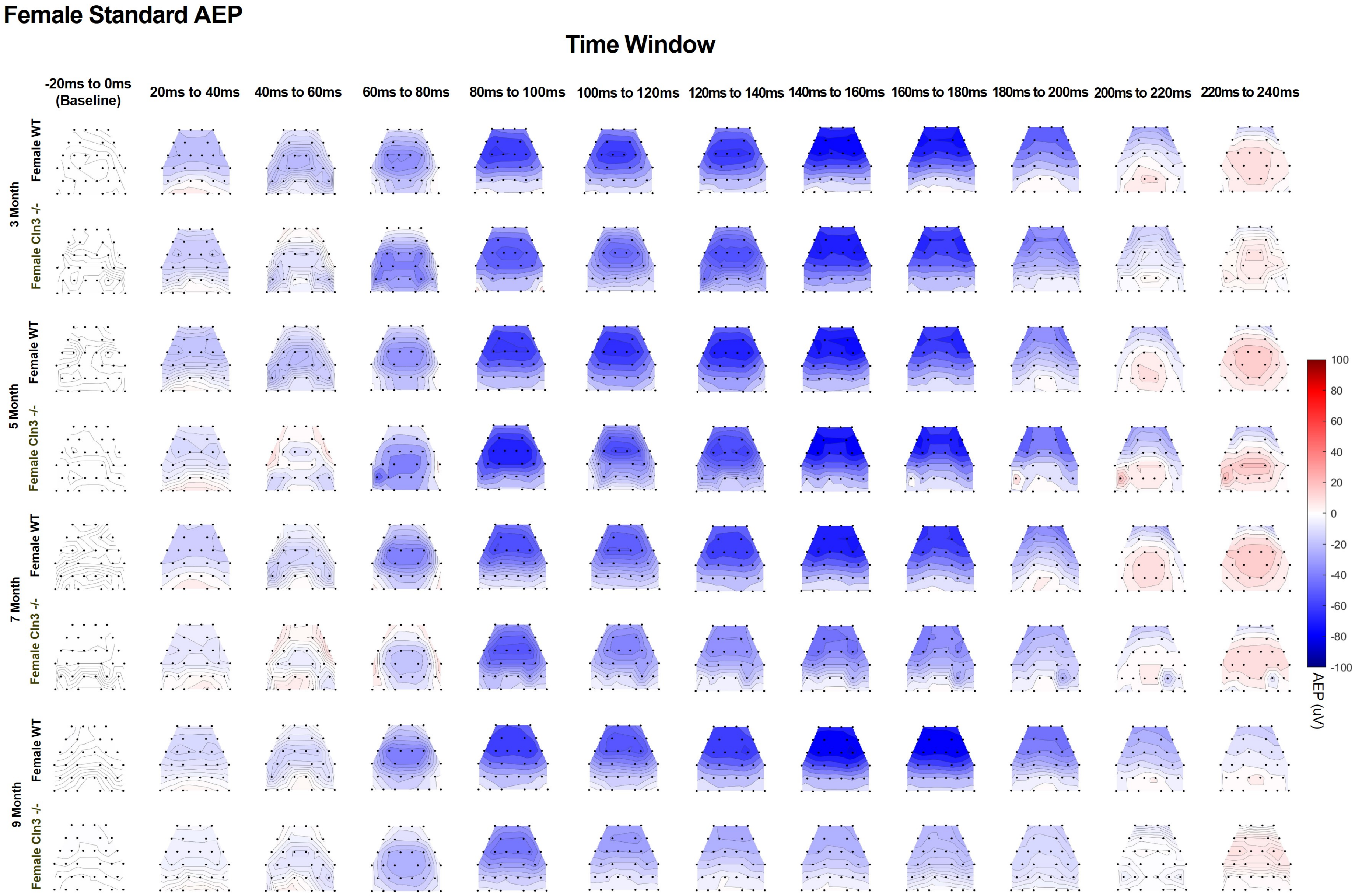
Female Standard AEP
Time Window
-20ms to 0ms (Baseline)
20ms to 40ms
40ms to 60ms
60ms to 80ms
80ms to 100ms
100ms to 120ms
120ms to 140ms
140ms to 160ms
160ms to 180ms
180ms to 200ms
200ms to 220ms
220ms to 240ms
3 Month
5 Month
7 Month
9 Month
Female WT
Female Cln3 -/-
100
80
60
40
20
0
-20
-40
-60
-80
-100
AEP (uV)

**Extended Data Figure 7-1. Female *Cln3-/-* mice showed a progressive decline of standard AEPs in spatial distribution.** Averaged topographical maps of standard AEPs in a 20ms time window from female WT (n = 6 for all ages) and female *Cln3-/-* mice (n = 7 for all ages). Female *Cln3-/-* mice showed a progressive decline of standard AEPs in spatial distribution. Blue shows negativity and red shows positivity for AEP.

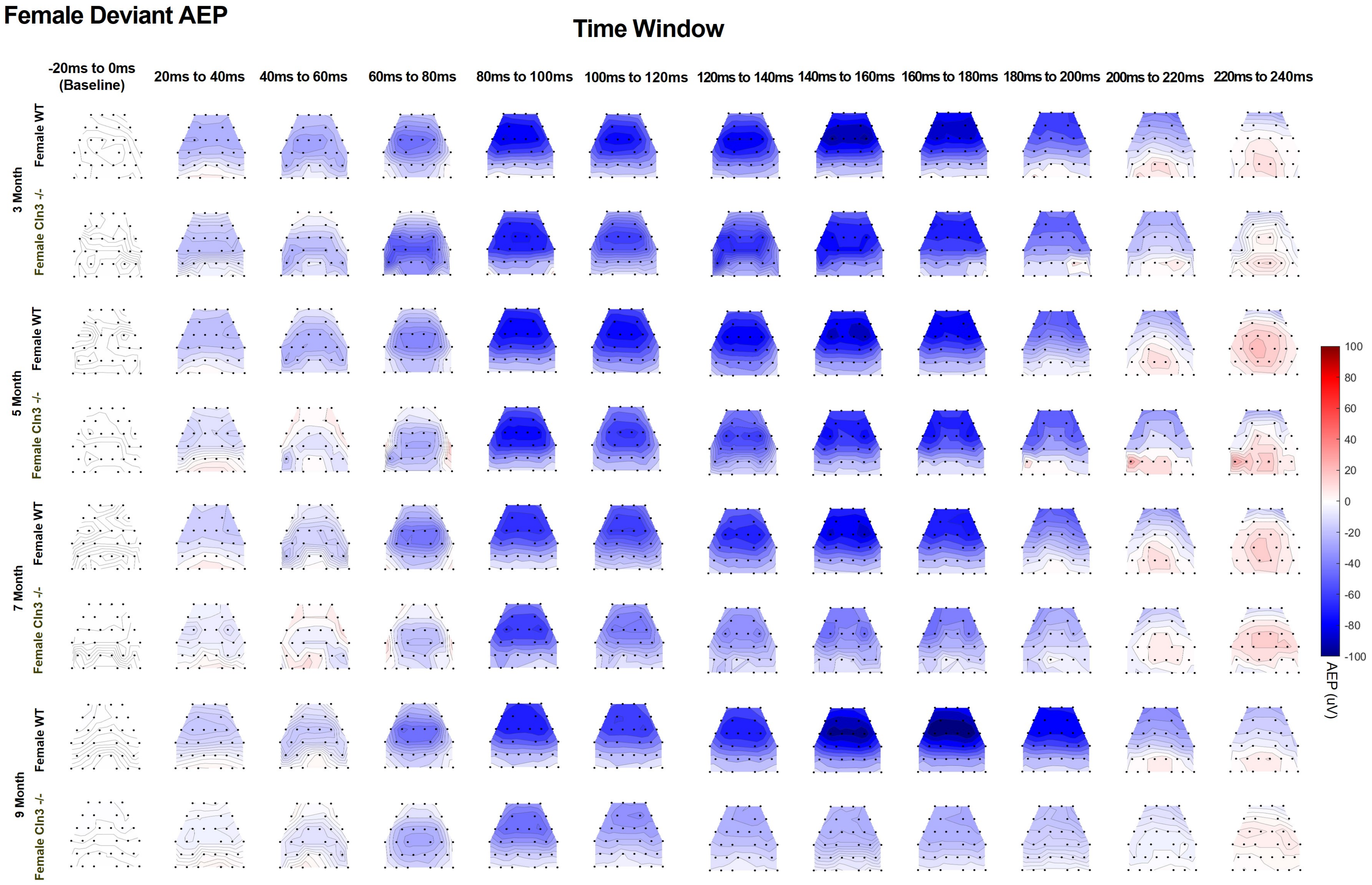

Female Deviant AEP
Time Window
-20ms to 0ms (Baseline)
20ms to 40ms
40ms to 60ms
60ms to 80ms
80ms to 100ms
100ms to 120ms
120ms to 140ms
140ms to 160ms
160ms to 180ms
180ms to 200ms
200ms to 220ms
220ms to 240ms
3 Month
5 Month
7 Month
9 Month
Female WT
Female Cln3 -/-
100
80
60
40
20
0
-20
-40
-60
-80
-100
AEP (uV)

**Extended Data Figure 7-2. Female *Cln3-/-* mice showed a progressive decline of deviant AEPs in spatial distribution.** Averaged topographical maps of deviant AEPs in a 20ms time window from female WT (n = 6 for all ages) and female *Cln3-/-* mice (n = 7 for all ages). Female *Cln3-/-* mice showed a progressive decline of deviant AEPs in spatial distribution. Blue shows negativity and red shows positivity for AEP.
